# Venomous Noodles: evolution of toxins in Nemertea through positive selection and gene duplication

**DOI:** 10.1101/2023.04.12.536482

**Authors:** G.G Sonoda, E.C. Tobaruela, J.L. Norenburg, J.P. Fabi, S.C.S. Andrade

**Affiliations:** Departamento de Genética e Biologia Evolutiva, IB-Universidade de São Paulo, São Paulo, Brazil; Laboratório de Toxinologia Aplicada, Instituto Butantan, São Paulo, Brazil; Faculdade de Ciências Farmacêuticas, Food Research Center (FoRC), Universidade de São Paulo, São Paulo, Brazil; Department of Invertebrate Zoology, Smithsonian’s National Museum of Natural History. Washington, DC 20560-0163

**Keywords:** Ribbon worms, dN/dS, Gene Duplication, Molecular evolution, Cytotoxin-A

## Abstract

Some, probably most and perhaps all, members of the phylum Nemertea are venomous, documented so far from marine and benthic specimens. Although the toxicity of these animals has been long known, systematic studies on characterization of toxins, mechanisms of toxicity and toxin evolution for this group are relatively scarce compared to other venomous groups. Here we present the first investigation of the molecular evolution of toxins in Nemertea. Using a proteo-transcriptomic approach, we described toxins in the body and poisonous mucus of the pilidiophoran *Lineus sanguineus* and the hoplonemertean *Nemertopsis pamelaroeae*. Using these new and publicly available transcriptomes, we investigated the molecular evolution of six selected toxin gene families. In addition, we have also characterized *in silico* the toxin genes found in the interstitial hoplonemertean, *Ototyphlonemertes erneba*, a first record for meiofaunal taxa. We identified 99 toxin transcripts in the pilidiophoran *L. sanguineus,* including previously known toxins, such as the *alpha-nemertides* and the *Cytotoxins-A*. However, for each of the hoplonemerteans, no more than 30 toxin transcripts were found. Genomic alignments and tree reconciliation methods supported the occurrence of at least one gene duplication in each analyzed toxin gene. Evidence of positive selection was observed in all investigated toxin genes. We hypothesized that an increased rate of gene duplications observed for Pilidiophora could be involved with the origin and expansion of toxin genes. Studies concerning the natural history of Nemertea are still needed to understand the evolution of their toxins. Nevertheless, our results show evolutionary mechanisms similar to those observed in other venomous groups.

## Introduction

Ribbon worm is the common name for animals belonging to the phylum Nemertea, a group with more than 1,300 valid species (Norenburg et al. 2023). The synapomorphy of the group is the presence of an eversible proboscis housed in a cavity called the rhynchocoel. With few exceptions, nemerteans are free-living predators or scavengers inhabiting the marine benthos (Brusca et al. 2016). Recently, it was proposed that the phylum should be divided into three classes: Hoplonemertea, Palaeonemertea, and Pilidiophora, the latter comprising the order Heteronemertea, and the family Hubrechtidae (Strand et al. 2019). Reciprocal monophyly of Pilidiophora, Hoplonemertea and Palaeonemertea was recovered in a phylogenomics approach by Andrade et al. (2014).

Although their soft body makes them look vulnerable and inoffensive, animals from this phylum possess toxins that may be used for predation and defense. The first known record of a ribbon worm was in 1555 (Göransson et al. 2019) and includes the first description of its venomous nature. However, more detailed and systematic studies of the toxin molecules was not published until the 1930s (Bacq 1936; Bacq 1937). Since then, more studies have addressed the biochemical activity of these toxins (Kem et al. 1971; Kem 1976; Kem and Blumenthal 1978; Kem 1985; Berne et al. 2003; Jacobsson et al. 2018) and their role in the natural history of Nemertea (McDermott 1976; Thiel and Kruse 2001).

Toxins are defined as substances produced by living organisms that cause deleterious effects in other organisms (named targets) exposed to them (Wexler et al. 2015). Due to their potential to act as pharmacological agents and pesticides, the isolation and description of these substances are common practices. The ecological role of the toxins in Nemertea has been largely deduced from field and laboratory observations coupled with biochemical assays (McDermott 1984; Kem 1985; Christy et al. 1997; Thiel and Kruse 2001). It is known that the proboscis plays a fundamental role in aiding the delivery of these toxins. Hoplonemerteans bear stylets on the proboscis that pierce a target’s body wall, creating a wound through which toxin-containing substance enters the target organism (McDermott 1976; McDermott and Roe 1985). In the case of Monostiliferous hoplonemerteans, toxin may be actively introduced via the pumping action of a specialized, muscular, mid-proboscis bulb (Stricker, 1985). Palaeonemerteans and pilidiophorans lack any rigid piercing structure. These worms rely on the physical contact of the proboscis coated in a poisonous mucus to envenom their prey (McDermott and Roe 1985). In addition, toxins can be found in the mucus surrounding the animal’s body, acting as a potential defense mechanism against predators. Such mucus was demonstrated to contain peptides with neurotoxic and cytotoxic properties (Posner and Kem 1978; Berne et al. 2003; Jacobsson et al. 2018).

Both proteinaceous (Kem 1976; Kem and Blumenthal 1978; Berne et al. 2003) and non-proteinaceous (Kem 1971; Kem et al. 1976; Kem 1988; Asakawa et al. 2013) toxic compounds have been isolated from the mucus of different nemertean species. There is evidence that some of these toxins have potential application as pesticide (Jacobsson et al. 2018).

The evolution of proteinaceous toxins can be assessed and interpreted using molecular approaches. So far, the only Nemertean proteinaceous toxins that have been successfully isolated and had their toxicity assayed were obtained from Pilidiophora specimens. However, Whelan et al. (2014), found putative toxins in transcriptomes of the three classes of Nemertea. Recently, proteomic and transcriptomic approaches showed new potential proteinaceous toxins in the mucus and proboscis of hoplonemerteans (von Reumont et al. 2020; Verdes et al. 2022). These findings point toward various unexplored proteinaceous toxins in Nemertea, with bioactivity potential yet to be investigated.

Despite this tantalizing potential of Nemertea as a hotbed of toxin evolution, the phylum has received little attention in reviews of metazoan toxins (Whelan et al. 2014), while few groups, such as scorpions, snakes, spiders, and cone-snails have been in the spotlight for studies concerning biology and evolution of venoms (von Reumont et al. 2014). These studies point to clear patterns in the evolution of toxin genes and their origins. The classical and main hypothesis for their origin is through the duplication of both endogenous and toxin genes, which may go under neofunctionalization (Kordiš and Gubenšek 2000; Casewell et al. 2013). The multilocus toxin gene families support this hypothesis in cone-snails (Duda Jr and Palumbi 1999; Pi et al. 2006; Duda Jr and Remigio 2008; Chang and Duda Jr 2012), spiders (Binford et al. 2009), snakes (Fry et al. 2003; Juárez et al. 2008; Bayona-Serrano et al. 2020; Giorgianni et al. 2020) and scorpions (Ma et al. 2012; Zhu et al. 2012; Sunagar et al. 2013), many of which include endogenous genes (Fry et al. 2003; Sunagar et al. 2013; Bayona-Serrano et al. 2020; Giorgianni et al. 2020). Also, other mechanisms have been proposed to explain the origin of these genes, including alternative splicing (Cousin et al. 1998), modification of a physiological gene (Fry 2005), and lateral gene transfer (Cordes and Binford 2006). The multilocus toxin gene families also indicate that further gene duplication events are common in the evolutionary history of toxins. This might result from diversification of the chemical arsenal of organisms by neofunctionalization, thereby increasing the number of potential molecular targets for a given venom, which may increase their fitness.

Another critical pattern identified in the evolution of toxins is the adaptive (or positive) selection in toxin gene families, inferred through the non-synonymous/synonymous mutation ratio (dN/dS), as observed in toxin genes of cone-snails (Duda Jr and Palumbi 1999; Chang and Duda Jr 2012), scorpions (Zhu et al. 2012; Zhang et al. 2015), snakes (Nakashima et al. 1993; Juárez et al. 2008) and Spiders (Binford et al. 2009). As described by Van Valen’s Red Queen hypothesis (Van Valen 1977), a coevolution process between producer and target might trigger this particular pattern (Casewell et al. 2013). Positive selection in these gene families could also be a by-product of gene duplications followed by neofunctionalization, as the new gene copy product may evolve into a protein with an adaptive role (Han et al. 2009).

The main goal of this study was to investigate the molecular processes behind toxin evolution in Nemertea and to search for patterns described in the evolution of toxins in other metazoans. By investigating toxin evolution in this poorly studied Phylum and expanding the number of taxa investigated so far, our results will contribute to a better and less biased understanding of the origins and forces that drive the evolution of toxin genes in metazoans. To that end, we have sequenced transcriptomes of three Nemertea species, the cosmopolitan heteronemertean *Lineus sanguineus* (Rathke 1799), and the hoplonemerteans *Ototyphlonemertes erneba* Corrêa 1950 and *Nemertopsis pamelaroeae* Mendes et al. 2021. In addition, proteomic approaches were used to validate the new putative proteinaceous toxin found. The use of multi-omic approaches and the availability of previously sequenced data from the three Nemertea classes made it possible to analyze this molecular evolution in Nemertea from an integrative perspective with an emphasis on Pilidiophora, shedding light on the evolutionary scenario of the toxins of an understudied and ecologically crucial group.

## Material and Methods

### Sampling

Specimens of *Lineus sanguineus, Nemertopsis pamelaroeae* and *N. berthalutzae* Mendes et al. 2021 were collected in beds of the oyster *Crassostrea sp.* along the Southeast Brazilian coast (Table 1). Large portions of the beds were collected using a hammer and a putty knife. The fragments were placed in trays containing seawater at room temperature, left for two hours until the individuals crawled to the water surface, and then removed with a thin brush. *Ototyphlonemertes erneba* specimens were collected by dripping water in a tray with collected sediment, following Corrêa’s protocol (Corrêa 1949). The species were morphologically identified using Envall and Norenburg (2001) key characteristics list. All samples were preserved in RNAlater (Table 1). Animals were sampled under permits issued by Institute Chico Mendes (ICMBio), protocol numbers 55701 and 67004.

**Table 1:**
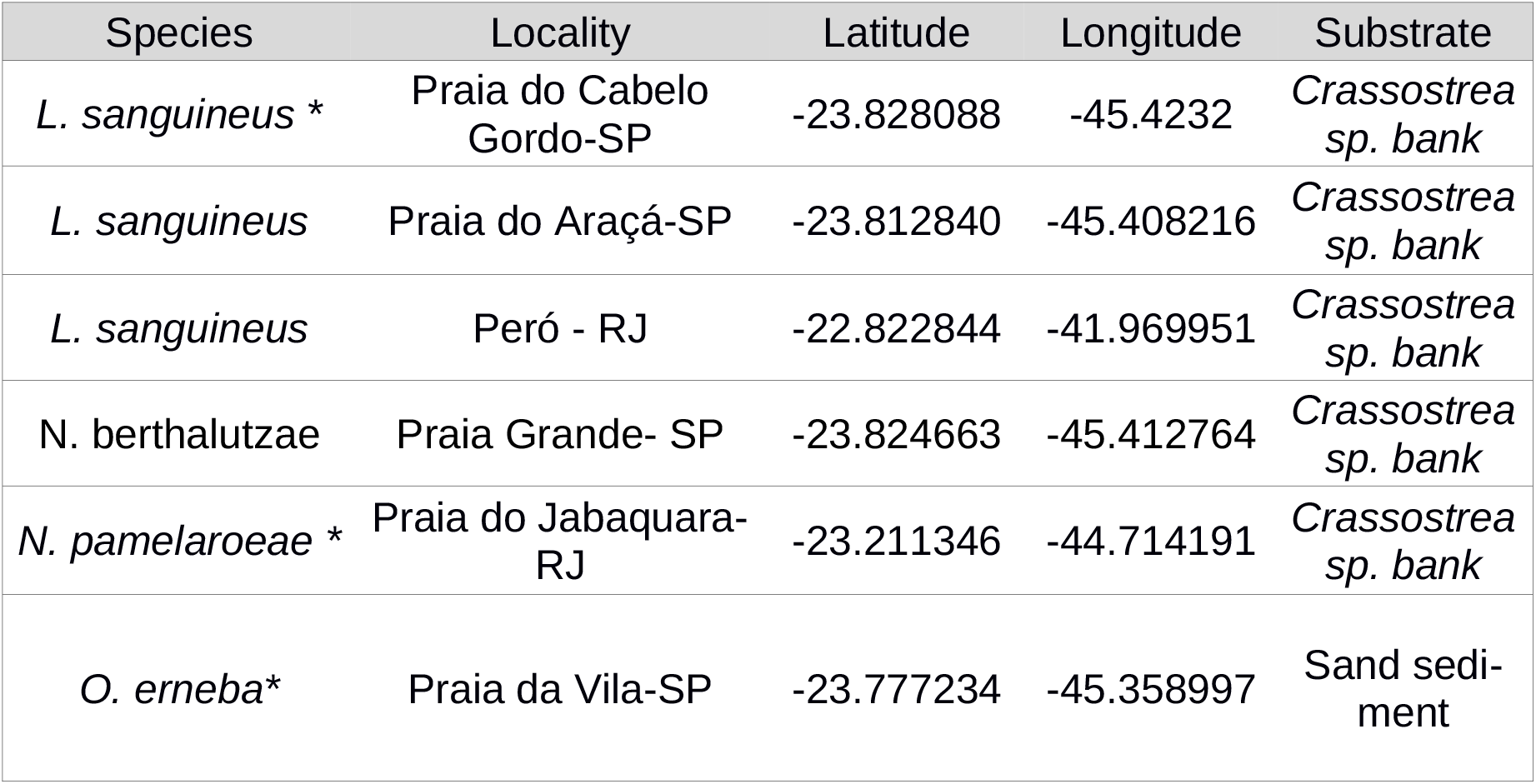
Species collected and their respective collection local and substrate. * samples used for RNA-Seq, the remaining were used for proteomic.

### RNA extraction and Sequencing

Total RNA was extracted using Tri-Reagent (Ambion), using the RNA extraction and purification protocols described in Riesgo et al. (2012).

For *Nemertopsis pamelaroeae* and *Lineus sanguineus* extractions, the whole animal was used. For *Ototyphlonemertes erneba* transcriptomes, extraction was performed using a pool of 12 individuals, since a single sample did not yield enough RNA. The RNA quality and quantity was assessed in a Qubit fluorometer v.3 (Thermo Fisher Scientific, Waltham, MA, USA), and samples with ratios between 1.8 - 2.2 were considered pure. Sample integrity was confirmed with BioAnalyser equipment (Agilent Technologies Inc., Santa Clara, CA, USA). A cDNA library was produced from each sample using the TruSeq RNA Library Preparation V2 kit (Illumina Inc., San Diego, California, USA). Libraries were clustered and sequenced on the Illumina HiSeq 2500 platform with 2 × 100 bp paired-end, using a TruSeq SBS V3 kit (Illumina Inc., Thermo Fisher Scientific), at the Laboratório de Biotecnologia Animal, Escola Superior de Agricultura “Luiz de Queiroz” – Universidade de São Paulo (ESALQ-USP).

Raw reads from 21 Nemertea taxa comprising 5 Palaeonemertea, 10 Pilidiophora and 6 Hoplonemertea species transcriptomes, were downloaded from the NCBI database using the SRA toolkit (Sherry et al. 2012) and are presented in Table S1. These were selected based on the availability of paired-end Illumina reads from the body transcriptome. Two additional *Lineus sanguineus* transcriptomes were included. Transcriptomes from all individuals were used for phylogeny inference, gene duplication analysis, and selection tests (Figure S1). Assembly completeness was evaluated using the database metazoa_odb9 from BUSCO (Waterhouse et al. 2018).

### Transcriptome assembly and ORF prediction

Prior to the assembling, the overall quality of the reads was evaluated using the software FastQC v0.11.3 (Andrews 2010). Reads with average quality under 24 phred score, adapters and putative contaminant reads were filtered out using the software Seqyclean v.1.10.09 (Zhbannikov et al. 2017) and the UniVec (https://www.ncbi.nlm.nih.gov/tools/vecscreen/univec/) as putative contaminant database. The transcriptomes were assembled using the software Trinity v2.8.4 (Grabherr et al. 2011). TransDecoder v5.5.0 (Haas and Papanicolaou 2016) was used to predict the translated proteins and their respective coding sequences (CDS), with a cut-off of a minimum of 30 amino acids for prospection of toxin genes, selection tests and duplication analysis of toxin genes; and cut-off of 100 amino acids for the phylogenomic analysis and overall gene duplication analysis (Figure S1). The 30 amino acid cut-off was defined considering the previous report of toxins smaller than 100 amino acids, such as the alpha and beta-nemertides (Jacobsson *et* al. 2018). While the higher cut-off for the phylogenomics and overall gene duplication analysis was chosen to avoid orthogroups with short sequences, potential mis-assemblies and false open reading frames (ORFs).

### Phylogenomic analysis

To proceed with the phylogenomic and molecular evolutionary analysis, predicted proteins for transcriptomes of each species were assigned to orthogroups using the software OrthoFinder v2.3.3 (Emms and Kelly 2015) with default parameters.

The orthogroups were filtered according to their occupancy, and only groups with at least 26 taxa (∼80% occupancy) were kept. Each orthogroup was aligned using MAFFT L-INS-i v.7.407 (Katoh et al. 2005), and each gene tree was obtained using IQTree v.1.6.12 (Nguyen et al. 2015). Node support was evaluated with 100 ultrafast bootstrap replicates. FASTA-formatted files were trimmed with TrimAl v1.2 to account for alignment uncertainty, with a gap threshold of 80% and conserving a minimum of 20% of the original alignment (Capella-Gutiérrez et al. 2009). Monophyly masking was performed by an iterative paralogy pruning procedure using PhyloTreePruner (Kocot et al. 2013).

The package BaCoCa v.1.105r (Kück and Struck 2014) was used to estimate the Relative Composition Frequency Variability (RCFV), which measures the absolute deviation from the mean for each amino acid and for each taxon and sums these up over all amino acids and all taxa (Zhong et al. 2011). Based on the values distribution, partitions with a high degree of compositional heterogeneity were filtered out.

The concatenated matrix was analyzed by IQ-Tree, using LG4X+G as evolution model and 1,000 bootstrap replicates. Representatives of the phyla Annelida and Mollusca were chosen as outgroups, based on previous studies on metazoa relationships (Laumer et al. 2019). The species chosen as outgroups were the Annelida *Phyllochaetopterus* sp., *Myzostoma seymourcollegiorum,* and *Capitella teleta*; and the Mollusca *Solemya velum*, *Lottia gigantea*, and *Monodonta labio* (Table S1).

### Toxin identification

The transcriptomes of the three sequenced species (the pilidiophoran *Lineus sanguineus;* and the hoplonemerteans *Nemertopsis pamelaroeae and Ototyphlonemertes erneba)* were assessed for transcripts encoding toxins. Their predicted proteomes (see *Transcriptome assembly and ORF prediction* section; Figure S1) were annotated using BLAST V2.9.0 (Altschul et al. 1990) to identify the proteins in the Swiss-Prot database (The UniProt Consortium et al. 2021) to produce the best alignment for each predicted protein, with a maximum e-value of 10e-4. Predicted proteins with the best hits containing the Gene Ontology term “Toxin activity; GO:0090729” were considered putative toxins. Additionally, as we expect a high abundance of toxins in the mucus proteome, we annotated the proteins detected in mucus samples (see below) with more permissive parameters (e-value < 10). We manually reviewed the annotation of the proteins found in the mucus to identify additional putative toxins. Identified putative toxins were further annotated against the same database using phammer (HMMER suite v3.3, Mistry et al. 2013). Finally, their domains were predicted using hmmscan, also from HMMER suite, and the Pfam 33.1 database (Mistry et al. 2021).

As this study focused on the toxins from Nemertea, we filtered out prokaryotic toxins. Also, transcripts annotated as Perivitelline 2 were filtered out. Although these proteins are annotated with the GO term Toxin activity, they are mostly known for their role in the nutrition and defense of eggs (Dreon et al. 2013, although see Cadierno et al. 2017).

### Proteomic of mucus and whole individuals

The presence of the predicted proteins in either the body or the mucus proteome validated the transcriptomic results. Also, we considered the presence of the predicted putative toxins in the mucus as evidence for its role as a toxin. We performed proteomic experiments with four *Lineus sanguineus* and two *Nemertopsis berthalutzae,* also collected in Southeastern Brazil (Table 1). *N. berthalutzae* was used as no new fresh samples from *N. pamelaroeae* were found in the fieldwork. As sister species to *N. pamelaroeae*, both belonging to the *Nemertopsis bivittata* complex (Mendes et al. 2021), we expected similar defensive toxins between these species, which coexist on the coast and occupy the same habitat (Mendes et al. 2021), meaning they might face the same predators and feed on the same prey. Before mucus extraction, a sample of the water in which animals from each species rested for 10 minutes was collected as a control. After that, animals were stimulated to secrete mucus via repeated aspiration and expiration from a plastic pipette and manual shaking of a 2 mL vial containing approximately 1.5 mL of seawater. Three cycles of ten aspirations followed by 10 seconds of manual shaking were performed for each animal, with a 10-minute interval between each cycle. Animals were also carefully scraped with a hypodermic needle to remove the additional mucus from the skin. Afterward, the samples, the seawater containing the mucus and the control were separately stored in liquid nitrogen, then lyophilized and stored at -20° C (protocol adapted from H. Andersson, personal communication).

Lyophilized mucus samples were resuspended in milli-Q water, and excess salt was removed by size exclusion chromatography using the gravity protocol of columns packed with Sephadex^®^ G-25 Medium (PD-10, Cytiva) as in Jacobsson et al. (2018). The desalted samples were lyophilized again before dithiothreitol (DTT) reduction, iodoacetamide (IAA) alkylation, and digestion with Trypsin and Lys-C (Trypsin/Lys-C Mix, Mass Spec Grade Promega) for 14 hours at 37° C.

For the body proteome, individuals with dry mass ranging between 1.1 mg and 3.5 mg were manually homogenized with micropistilles in a solution of LAEMMLI with 10% beta-mercaptoethanol. The final concentration was 7.2 μg of animal/μL. In order to avoid the more abundant structural proteins overshadow the toxins, samples were fractionated in an 12.5% SDS-PAGE following Green and Sambrook (2012). The resulting gel was cut in three fragments accordingly to protein content visualized by Coomassie blue G staining and were submitted to in-gel digestion. The used protocol was a modified version of Shevchenko et al. (2006), in which proteins were reduced (DTT) and alkylated (IAA) before the digestion with Trypsin and Lys-C (Trypsin/Lys-C Mix, Mass Spec Grade Promega) for 14 hours at 37° C. Lastly, the digested peptides of both mucus and bodies were desalted using C18 ZipTips columns (Millipore) and concentrated using a CentriVap (Labconco Corporation).

Peptides were redissolved in 0.1% formic acid (FA), injected in a C18 pre-column (Acclaim PepMap RSLC Nano-Trap column; 3 μm, 100 Å, 75 μm × 20 mm, Thermo Fisher Scientific) and separated on a C18 analytical column (Acclaim PepMap RSLC column; 2 μm, 100 Å, 75 μm × 150 mm, Thermo Fisher Scientific) using a linear gradient from 5% buffer A (80% acetonitrile, 0.1% FA) to 95% for 120 min, with buffer B containing 0.1% FA. The flow rate was set to 350 nL/min and oven temperature to 50° C. The QToF Impact II Bruker mass spectrometer (Bruker Daltonics, Bremen, Germany) was interfaced to the nanoLC system (Thermo Scientific UltiMate™ 3000 RSLCnano system) with the CaptiveSpray nanoBooster source (Bruker Daltonics) using acetonitrile as dopant. Liquid Chromatography coupled to mass spectrometry data were acquired using a Data Dependent Acquisition (DDA) scheme. Briefly, a MS scan at scan speed of 200 ms was followed by 20 MS/MS fragment scans of the most intense precursors (50 ms) in a total cycle time of 1.2 seconds. The mass range of the MS scan was set from *m/z* 150 to 2200. Isolation of precursor ions was performed using an *m/z* isolation window of 2.0 and a dynamic exclusion of 0.4 min. The collision energy was adjusted between 23 and 65 eV as a function of the *m/z* value. All samples were analyzed in random order.

PEAKS Studio Xpro (V10.6 PEAKS Team) was used to query scans against proteomes containing proteins with 30 amino acids or more (Figure S1), predicted from the transcriptomes of *Lineus sanguineus* or *Nemertopsis pamelaroeae*, in addition to the SwissProt dataset of proteins to search for contamination. Precursor mass search was set to monoisotopic, parent mass error tolerance was set to 25 ppm, fragment mass error tolerance was set 0.1 Da and maximum FDR was set to 5%. The enzyme was set to Trypsin/Lys-C, and the maximum missed cleavage sites were set to three. For the mucus analyses, we considered peptides as contaminants if their peptides area in the seawater sample was more significant than in the mucus sample.

### Trees and alignments for toxin orthogroups

We only considered putative toxins validated in the proteomes for the evolutionary analysis. From these, we selected six putative toxins of *Lineus sanguineus*, based on their presence in the transcriptomes of other species and on previous reports of these toxins.

Orthogroups containing these toxins and their homologous genes were selected. The translated amino acid sequences of these orthogroups were replaced by their respective CDS, previously inferred with TransDecoder. Redundant CDs within each orthogroup were filtered out using cd-hit-est v4.8.1 (Fu et al. 2012), with 100% similarity. The remaining sequences were aligned by codon with Muscle at MEGA-X (Kumar et al. 2018). We manually removed highly divergent and incomplete sequences for toxins found in the mucus. As the orthogroups containing the toxins found in the body had more sequences, MaxAlign v1.1 (Gouveia-Oliveira et al. 2007) was used to automatically remove sequences while keeping maximum gap-free alignment area. Maximum likelihood trees for these alignments were determined using RAxML v8.2.12 (Stamatakis 2014) rapid bootstrap analysis (N=100, model = GTRGAMMA). The resulting trees were midpoint rooted.

### Gene duplications in toxin orthogroups

To detect gene duplications, all gene trees were compared to the phylogenomic species tree. We assumed that orthogroups determined by OrthoFinder contain predicted proteins that share homology, including orthologs and paralogs (Emms and Kelly 2015). It implies that each sampled transcriptome containing more than one transcript in the orthogroup has experienced gene duplication. To detect gene duplications, gene tree reconciliation analyses were performed using DLCpar v2.0.1 (Wu et al. 2014), which uses dynamic programming to find the most parsimonious set of gene duplications, losses, and deep coalescence events that explains the observed data. As this analysis is based on transcriptomic data, it is not possible to distinguish gene duplications in one species from isoforms derived from alternative splicing of the same gene. So, duplication events leading to two sequences of the same species were counted out. Also, transcriptomic data does not allow for distinguishing between gene losses and the absence of transcription during sample fixation. Therefore, the gene loss results were not accounted for it.

Since transcriptomic data can lead to a false interpretation of the number of genetic copies due to post-transcriptional variations, such as splicing, we validated these results using a combination of genomic and transcriptomic data. For such, *Lineus longissimus* transcripts (Table S1) contained in toxin orthogroups were aligned to the genome of *L. longissimus* (PRJEB45185; Kwiatkowski et al. 2021) using blat v. 36 (Kent 2002) with default parameters. Alignments with introns bigger than 20kb were filtered out. Alignments were visualized with IGV-Integrative Genomics Viewer v2.14.1 (Robinson et al. 2011) and the R package GVIZ (Hahne and Ivanek, 2016). We defined the gene copy number as the different genomic regions where the transcripts aligned.

### Nemertea gene duplication events

Overall estimates of gene duplication were performed using OrthoFinder, which infers gene duplications by comparing a species tree given by the user (in this case, the phylogenomic tree) with rooted gene trees inferred using the OrthoFinder ‘msa’ algorithm with default options. Such comparison is done through a hybrid algorithm of DLCpar and species-overlap (Emms and Kelly 2015). For this analysis, we used the predicted proteomes with proteins containing 100 or more amino acids (Figure S1) to avoid false duplications inferred by mis-assembled small contigs and poorly predicted ORFs.

To further understand the expansion and contraction history of genes in the phylogeny of Nemertea, the transcriptomes were submitted to an estimation of birth-death rate of a given gene family per millions of years (ƛ), using CAFE, v4.2.1 (Han et al. 2013). To avoid misinterpretation of families with multiple isoforms, only the longest, and hence more informative, isoforms proteins (identified by Trinity) were included in the CAFE analysis. These passed through an all vs all blast and were clustered using MCL (Enright et al. 2002, inflation value=2.0). Clusters with 100 or more gene copies were filtered out to avoid gene families with high variance of copy numbers, as suggested by the author (Enright et al. 2002). An ultrametric tree was obtained with r8s from the phylogenomic tree following CAFE tutorial default protocol.

CAFE was used to estimate one scenario with one global ƛ value for the whole tree, another scenario allowing two different ƛ (one for Pilidiophora and another for the remaining taxa), and a third scenario with three ƛ (one for the *Lineus sanguineus* clade, one for the remaining Pilidiophora and one for the remaining taxa). This third scenario aimed to reduce any bias resulting from the inclusion of three individuals of the same species. The third scenario was also recalculated with an error model accounting for assembling errors. This model accounting for errors was finally used to estimate the expansion and contraction events in each node of the phylogenomic tree. To test if allowing more than one ƛ in the tree increased the models likelihood, twice the difference between the log likelihood of the global ƛ scenario and the multi ƛ scenario (2×(lnLikelihood_global_−lnLikelihood_multi_)), was compared to a null distribution of likelihood ratios (2×(lnLikelihood_global_−lnLikelihood_null_)) obtained by 194 null simulations (excluding -inf values) of the data set with a single ƛ for the whole tree, from which p-values were estimated (Han et al. 2013).

### Selection test

The orthogroups alignments obtained in the section “Trees and alignments for toxin orthogroups” and the gene trees reconciliation obtained in the section “Gene duplication in toxins” were tested for positive selection using ETE 3 (Huerta-Cepas et al. 2016), which employs CODEML, from the package PAML4 (Yang 2007). To test for positive selection in the whole orthogroup, we tested the alignment for positive selection using the site model. Also, subsets of three or more sequences identified as originating from a duplication event were marked for the branch-site selection test. By marking these subsets of sequences, we tested for positive selection in each subset defined by duplication events. The site model does not allow a more conserved portion of the sequence to mask sites being positively selected, which is more realistic than the branch model (Jeffares et al. 2015). The site model compares the near-neutral model M7, which assigns each site as either under negative selection (ω<1) or neutrality (ω = 1), with the model M8, which adds the possibility of sites being under positive selection (ω>1), to determine which one fits better the observed data. Similarly, the branch-site model tests if the data is better explained by a scenario that allows a marked set of nodes (foreground) to have sites under positive selection (bsA1) or a null scenario that restricts positive selection (bsA) in the whole tree.

## Results and Discussion

In this study, we investigated the molecular evolution of the toxins in the phylum Nemertea. Through a transcriptomic approach, using both published data and new transcriptomes we were able to identify known toxins and new putative toxins for Nemertea. Using a robust new phylogeny, we detected a higher gene duplication rate observed in Pilidiophora when compared to the remaining Nemertea classes, that could be associated with the origin of evolutionary novelties in this group (such as new toxin genes). In fact, we observed several events of gene duplication on the Pilidiophora toxins, a common feature in the evolution of toxin genes in other metazoans. In addition, evidence for positive selection was detected in six toxin genes, indicating an accelerated evolution comparable to toxin genes of more studied venomous taxa. These results are the first to empirically support that the evolution of toxins in Nemertea is similar to the evolution of toxins in most Metazoans.

### Transcriptomes assembly

After quality trimming, the raw Illumina reads were used to assemble the transcriptomes with TRINITY. The RIN and the main assembly metrics from *Lineus sanguineus, Nemertopsis pamelaroeae* and *Ototyphlonemertes erneba* are presented in Table S2. BUSCO analysis showed that the assembled transcriptomes presented homologs for a great majority of conserved metazoan genes dataset, indicating that the transcriptomes are highly diverse in terms of represented genes (Figure S2). The completeness of the downloaded transcriptomes ranged from 12% in *Tubulanus punctatus* to 97% in *Lineus viridis*. The low completeness from the *Lineus lacteus* is due to the fact that such data correspond to partial exome sequencing (Rousselle et al. 2020).

**Table 2:**
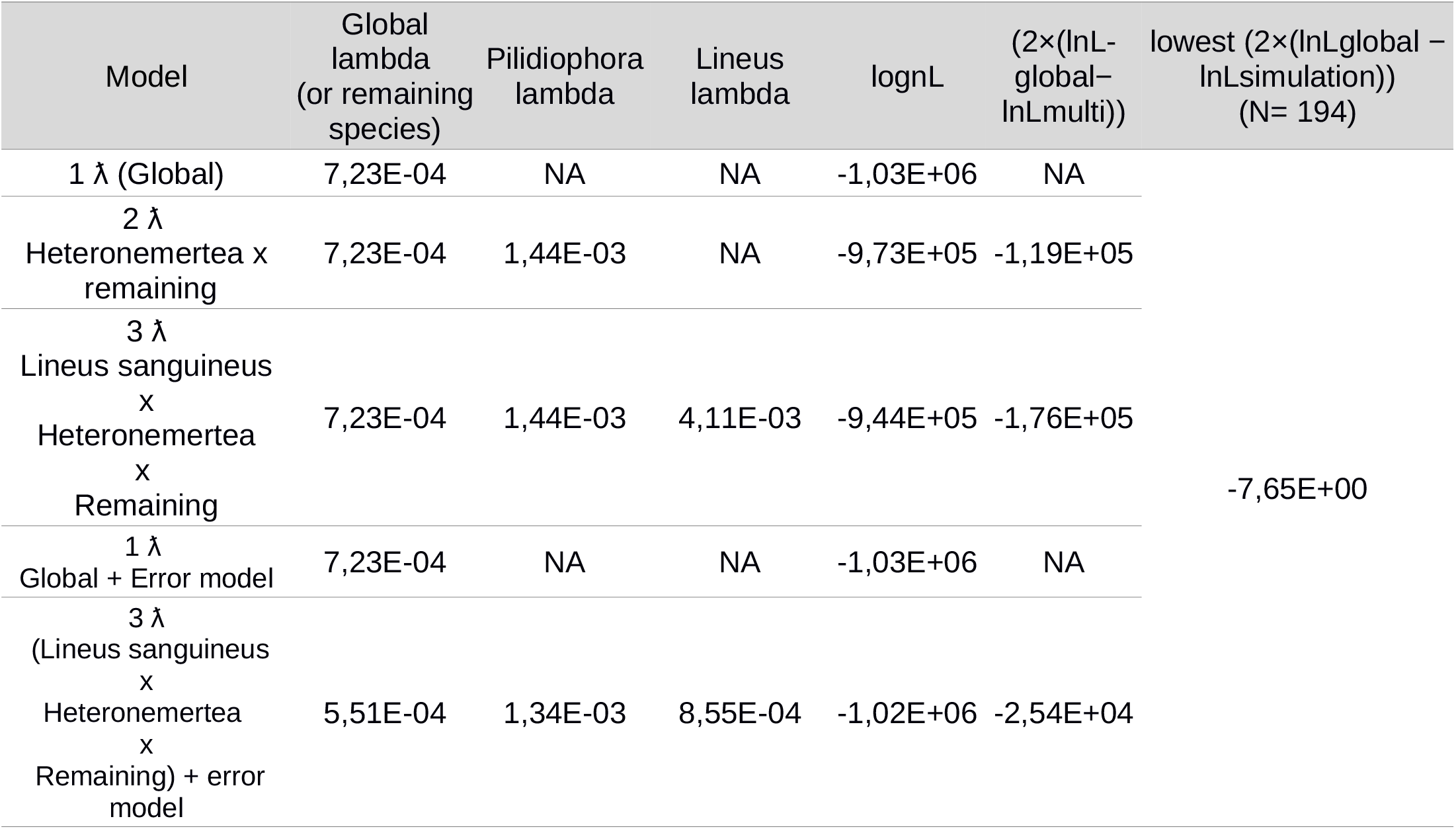
CAFE results. Lambda (estimate of gene duplication and contractions per gene family per unit of time) for each scenario. The formula (2×(lnLglobal−lnLmulti)) is used to test two different models by using their log likelihood. This value was compared to the values obtained in 194 null simulations. The number of Null simulations with lnL lower than the formula was the considered p-value. lognL: logn of likelihood; lnLglobal: logn of likelihood of single lambda model; lnLmulti: logn of likelihood of multi lambda models.

### Phylogenomic analysis

The assembled transcriptomes were used to infer a reliable species tree needed to comprehend better the history of gene duplications in toxin gene families. For such, proteomes predicted from the assembled transcriptomes were clustered with OrthoFinder. The number of represented orthogroups ranged from 1,815 in *Tubulanus punctatus* to 104,753 in the *Lineus sanguineus* collected in Brazil (Table S1). After occupancy filtering (Figure S3), the 5,208 orthogroups were submitted to monophyly masking. We evaluated the compositional heterogeneity effect on our 310,946 residuals aligned matrix. Based on the values distribution on a 95% confidence interval, the 271 orthogroups with the highest Relative Composition Frequency Variability (RCFV) were removed. The final matrix contained 299,880 aligned amino acids and 31 taxa.

The relationship of the major clades (Palaeonemertea, Pilidiophora and Hoplonemertea) remained as found in previous phylogenetic inferences based on genetic data (Figure 1) (Andrade et al. 2012; Andrade et al. 2014). Unlike Kvist et al. (2014), we did not recover a paraphyletic Palaeonemertea (Figure 1). The topology of both the Hoplonemertea and Heteronemertea groups are very similar to the ones found in literature, including the position of the collected taxa in the tree (i.e., *Lineus sanguineus, Nemertopsis pamelaroeae* and *Ototyphlonemertes erneba*) (Andrade et al. 2014; Kvist et al. 2014). The obtained phylogenomic tree suggests that *Riseriellus occultus* should be moved to the genus *Lineus*, which can also be observed in other works (Andrade et al. 2014; Kvist et al. 2014). All these findings are supported by nodes with 100% support value from the ultrafast bootstrap method. The specimen from the species *Lineus lacteus* clustered with the remaining *Lineus sanguineus*.

**Figure 1:**
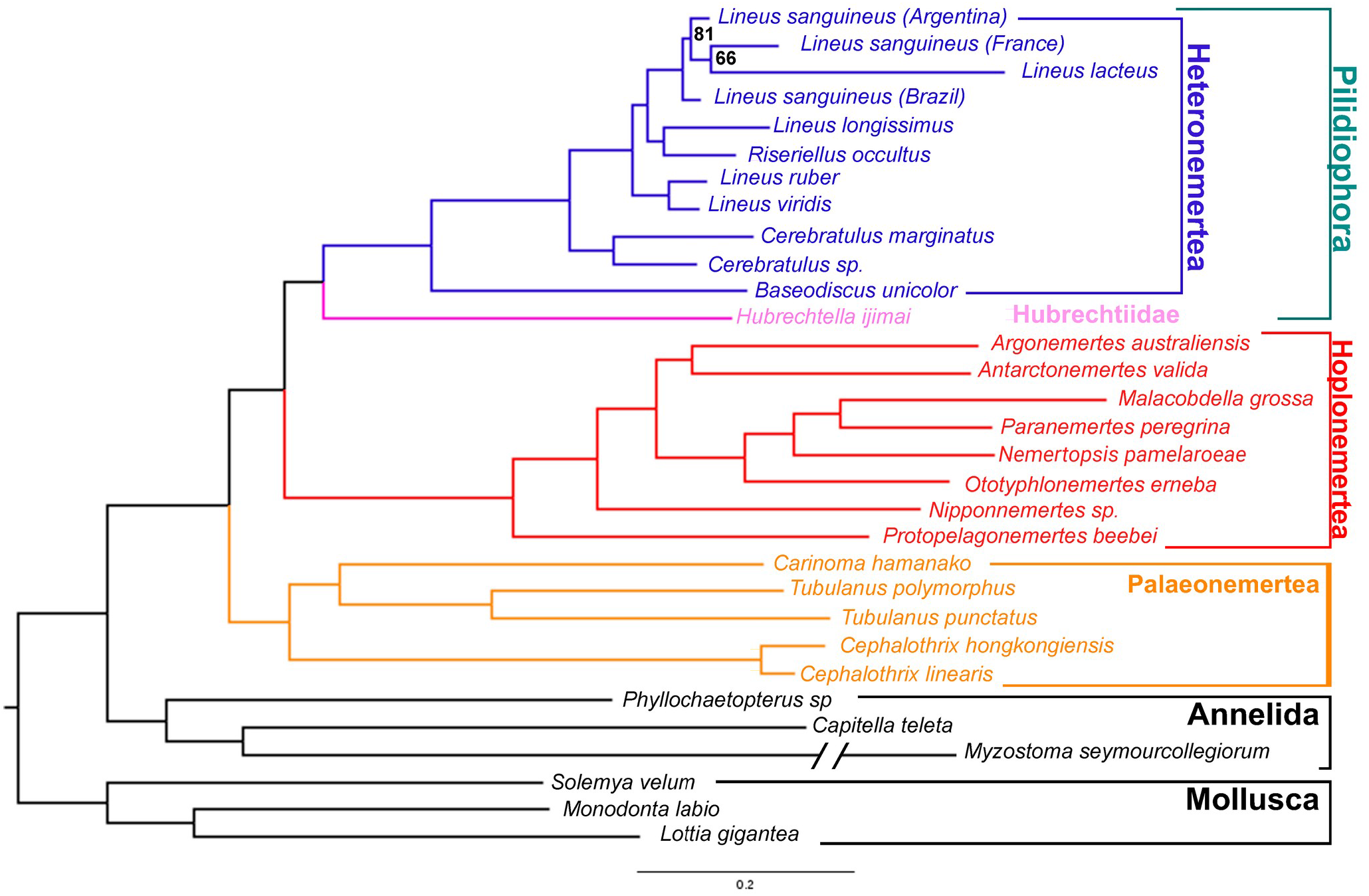
Phylogenomic tree of the Nemertea phylum obtained from 5208 genes (see methods). Tree was rooted to Mollusca. All the nodes have 100% bootstrap support except for the indicated.

### Identified putative toxins

The identification of putative toxins was based on sequence similarity to known toxins and their presence in the mucus proteome. Through the search by similarity, we found 85, 29 and 27 putative toxin transcripts for *Lineus sanguineus, Nemertopsis pamelaroeae,* and *Ototyphlonemertes erneba*, respectively. Moreover, we identified additional fourteen transcripts for *L. sanguineus* and one transcript for *N. pamelaroeae* as putative toxins for being present in the mucus proteomes and presenting features that indicate their role as toxins (Tables S3-S5). For the putative toxins found in the *L. sanguineus* mucus, these features include toxin domains such as cysteine knots (Pfam domain Toxin_35), PhTx neurotoxin family (De Figueiredo et al. 2001), Na/K-Atpase Interacting protein (Gorokhova et al. 2007) and Neurotoxin B-IV-like (Kem 1976) (Table S3). Regarding the putative toxin transcript found in the *N. berthalutzae* mucus, it presented similarity to a *metalloprotease 12A expressed in nematocyst* and to a toxin from *Amphiporus lactifloreus* (von Reumont et al. 2020), (Table S4; bitscore = 409; identities =35%; positives = 54%).

**Table 3:** Results from selection tests. *= Probability of site being under positive selection > 0.95; **= Probability of site being under positive selection > 0.99. NA: Not applicable, as these genes were not tested under the branch site model.

| Putative Toxin Gene | # of Sites under positive selection in Site-model | # of Sites under positive selection in branch site model | # of branches containing sites under positive selection (total branches tested) |
| --- | --- | --- | --- |
| *Cytotoxin-A* | 1** | 1* | 1 (3) |
| *Scoloptoxin-like* | 2*, 5** | 0 | 0 (2) |
| *Alpha-nemertide* | 1** | 1*, 1** | 1 (4) |
| *Ktx/myticin-like* | 1** | NA | NA |
| *Beta-nemertide* | 2** | NA | NA |
| *Gamma-Nemertide (agatoxin-like)* | 2* | NA | NA |

All these transcripts had a BLAST best hit in 52, 15 and 14 different proteins from the Swiss-Prot database, respectively for *L. sanguineus, N. pamelaroeae* and *O. erneba* (Table S3-S5; Figure S4). The best hits against the Swiss-Prot database include toxins from animals of different phyla, such as Cnidaria, Mollusca (genus *Conus*), vertebrates (snakes and fishes), arthropods (spiders and chilopods), and Echinodermata (crown-of-thorns starfish). This is the first report for many of these toxins in Nemertea. Whelan et al. (2014) found fewer toxin transcripts in transcriptomes of related species, probably due to differences in the toxin designation methodology.

The number of putative toxins found was considerably different between the pilidiophoran *Lineus sanguineus* and the two hoplonemerteans transcriptomes. Since we used the same toxin identification methodology for the three transcriptomes, and the BUSCO analysis showed a similar completeness level between them (Figure S2), this could indicate differences in the number of toxin genes expressed between the classes. Accordingly, Pilidiophora is the only nemertean clade from which proteinaceous toxins have been isolated and functionally characterized (Göransson et al. 2019). Even so, recent studies suggest a considerable diversity of proteinaceous toxins in Hoplonemertea (von Reumont et al. 2020; Verdes et al. 2022).

We can highlight the putative toxin transcripts identified as *Snaclec*, *Actitoxin* and the *U-nemertotoxin-1* (similar to *Plancitoxin-1*), found in the lineid and both hoplonemerteans putative toxin transcripts (Tables S3-S5); these were also found to be transcribed in the proboscis of *Amphiporus lactifloreus.* Both *Actitoxin* and *Plancitoxin-1* were also present in the defensive mucus proteome, meaning an actual role in predation and defense in that species (von Reumont et al. 2020). The finding of transcripts similar to these toxins in *Ototyphlonemertes erneba* (Table S5) is, to our knowledge, the first report of toxins from meiofaunal species. Although their toxicity is yet to be confirmed, their expression in the proboscis of a related species indicates that these also act as toxins in *O. erneba*. Furthermore, additional putative toxin transcripts such as *U-scoloptoxin(05)-Sm1a*, *Alpha-latrocrustotoxin* and a *Vulnericin* were identified solely from this meiofaunal species (Figure S4) and were unprecedented for Nemertea.

### Proteomic experiments

To confirm that the putative toxins identified in the transcriptome were translated and to identify more putative toxins, body and mucus proteomic analyses were made using high performance liquid chromatography with tandem mass spectrometry (HPLC-MS/MS). From the final set of 52 putative toxins from *Lineus sanguineus,* 16 were found in the mucus or body proteome, and 2 of 15 putative toxin transcripts from *N. pamelaoreae* were found in the mucus and body proteomes of *Nemertopsis berthalutzae*. Toxins identified in the transcriptome, but absent from the protein samples may have been obscured from the analyses by abundant structural and physiological proteins present in the body of both animals. Nevertheless, considering the small size of the animals, the low yeld of protein obtained in the mucus extraction and potential post-translational modifications, these results seem appropriate and provide more than enough data for subsequent analyses.

A total of 83,236 MS1 scans were obtained from all analyzed fractions of the four *Lineus sanguineus* bodies separately. The number of MS1 scans per fraction ranged from 13,837 and 13,951. From these scans, PEAKS Studio Xpro identified 2,577 proteins in the body, 12 of which were annotated as putative toxins (Figure S5a). For the *Nemertopsis berthalutzae* body, a total of 41,777 MS1 scans were obtained from fractions. The number of MS1 scans from fractions varied between 13,859 and 13,975. PEAKS Studio Xpro identified 576 proteins from the transcriptome of *Nemertopsis pamelaroeae* through these scans, two of which were considered toxins (Figure S5b).

The only toxin peptide with an area higher than 0 in the seawater control sample was an *Antistasin-like* peptide in *Lineus sanguineus* seawater, although Peaks did not find the same peptide in the mucus sample (Figure S6), which may indicate a potential misidentification exclusive to the seawater sample. For this reason, we did not consider any of the putative toxins as contaminants from the seawater. The mucus proteome from *Lineus sanguineus* and *Nemertopsis berthalutzae* yielded, respectively, 6,945 and 6,912 MS1 scans, while the control sample containing only seawater yielded 6,900 scans. PEAKS Studio Xpro assigned these scans to 25 and 82 different proteins derived from the studied species transcriptome, respectively for *L. sanguineus* and *N. berthalutzae*. Of these, 17 were identified as toxins in *L. sanguineus* (Table S3) and one was identified as a toxin in *N. berthalutzae* (Table S4).

The mucus proteome of the *L. sanguineus* and *N. berthalutzae* studies showed no sign of *Cytotoxin-A* or *U-nemertotoxin-1* homologs, contrasting with previous reports of toxins found in the mucus of *Parborlasia corrugatus* (Berne et al*. 2003)*, *Cerebratulus lacteus* (Kem and Blumenthal 1978) and *Amphiporus lactifloreus* (von Reumont et al. 2020). Since transcripts homologous to *Cytotoxin-A* and matching peptides were respectively found in the transcriptome and body proteome of *Lineus sanguineus* (Table S3), these toxins might have lost their defensive role and remain expressed only in the proboscis in this species, playing an important role in predation. Our results are the first to suggest different roles for these toxins in different species of Nemertea. Such differences could have evolved in response to diet changes and defense requirements (i.e., predators). Shifts in the composition of venom coupled with changes in its ecological role and delivery system are also present in spitting cobras, in which an increase in the cytotoxic activity of the venom is related to the defensive role of the spitted venom in causing pain (Kazandjian et al. 2021). Furthermore, snails from the genus *Conus* present distinct sets of toxins for defensive and predatory roles expressed in different parts of the venom duct (Dutertre et al. 2014). It can be compared to the different sets of toxins being expressed in the body tegument and proboscis tegument in Nemertea.

Other putative toxins found in the mucus of *L. sanguineus* were similar to *Antistasin*, a protein found in the salivary gland of leeches. *Antistasin* was found to be an anticoagulant and inhibitor of serine-proteases (Dunwiddie et al. 1989) Using MEME v.5.5.1 (Bailey et al. 2015), no anticoagulant motifs (obtained from Iwama et al. 2021) were found on the *Antistasin* sequences. In non-blood-feeding invertebrates, antistasin and proteins with similar domains are thought to have a role in immune responses, though their role remains unclear (Iwama et al. 2021). Verdes et al. (2022) found antistasin in the proboscis of the hoplonemertean *Antarctonemertes valida*, in which it likely is a predation toxin. Still, considering their capability of inhibiting serine-proteases, which often plays a digestive role in marine organisms (Barzkar et al. 2021), together with the the lack of anticoagulant motifs in these sequences, one can speculate that the presence of antistasin in the defensive mucus of *Lineus sanguineus* might inhibit the digestion in an eventual predator, rather than interfere with its hemostasis. In accordance with that, it was observed that the anemone *Metridium senile* kept in aquariums regurgitates ingested *Cerebratulus lacteus* (J. Norenburg, personal observations).

### Selected toxin gene families

Based on previous report of toxicity and presence in proteome, the following putative toxin genes were selected for further analysis: *Cytotoxin-A*, *scoloptoxin SD976-like*, *alpha-nemertide*, *beta-nemertide*, *U2-Agatoxin-like* and a putative toxin showing low similarity to *alpha-Ktx* (blast bitscore = 28.5)*, beta-defensin* (phmmer score = 23.2) and *myticin* (Pfam score = 16). Transcripts from *scoloptoxin SD976-like* were found in both *L. sanguineus* and *N. pamelaroeae* transcriptomes, while transcripts from the remaining toxins were exclusive to *L. sanguineus*. Peptides derived from the *scoloptoxin SD976-like* were found in the body proteome of both *L. sanguineus* and *N. berthalutzae*. In contrast, peptides from C*ytotoxin-A* were only found in the body proteome of *L. sanguineus* and the remaining were only found in the mucus proteome of *L. sanguineus* (Table S3-S4; Figure S5).

Final codon-alignments for the *Cytotoxin-A*, *scoloptoxin-SD976-like*, *alpha-nemertide*, *beta-nemertide*, U2-Agatoxin-like and *alpha-KTx-like* had, respectively, 598, 2058, 342, 273, 351, and 204, base pairs (Figures 2, S7-S11). These alignments and derived gene trees were used to perform selection tests and infer gene duplications by an algorithm of tree reconciliation.

**Figure 2:**
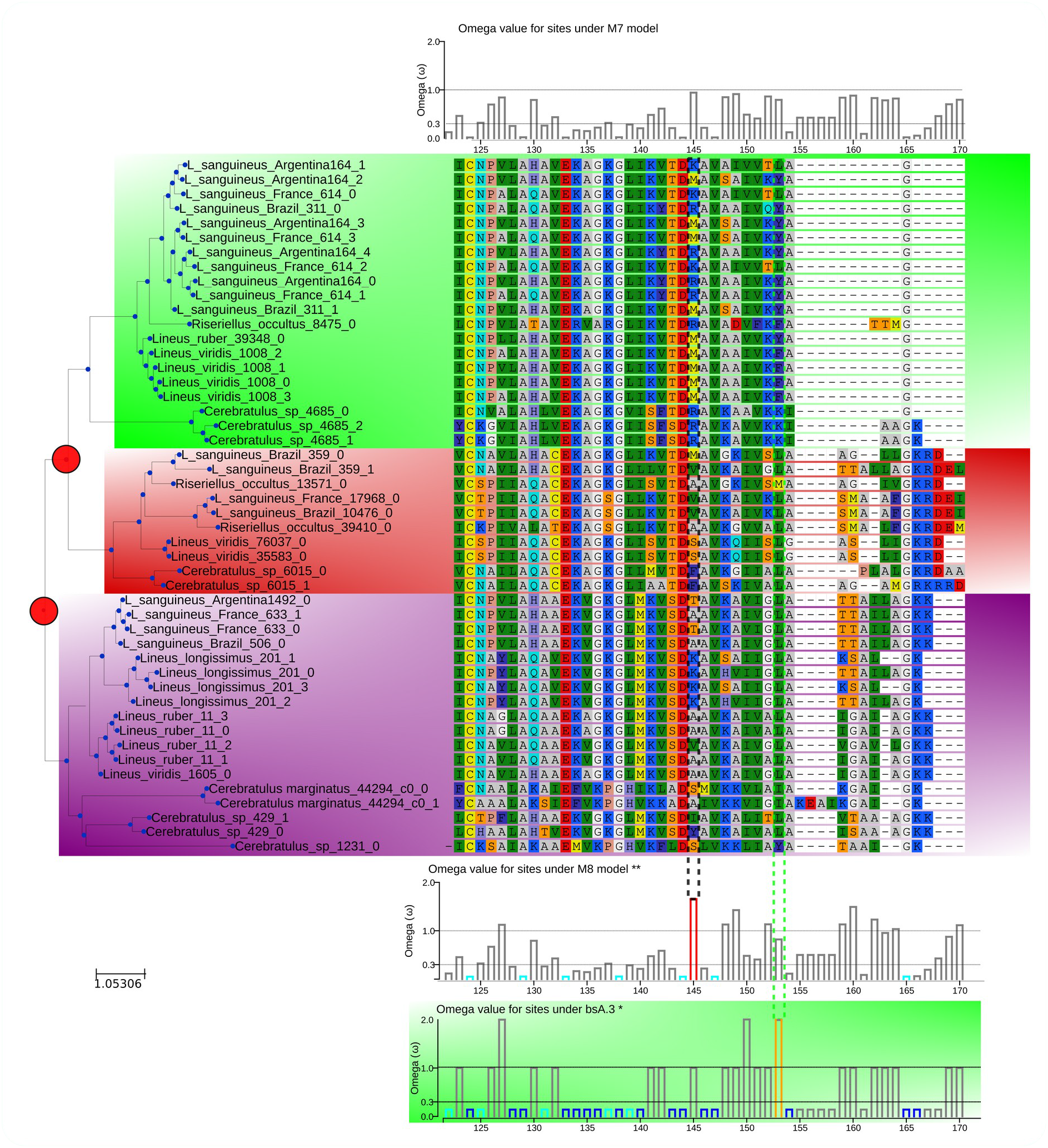
Gene tree reconciliation, alignment and omega value for sites of the Cytotoxin-A Orthogroup. The reconciled gene tree (left) was inferred by comparing the maximum-likelihood gene tree and the species tree using DLCpar to identify the most parsimonious history of gene duplications (red circles in the tree), losses and deep coalescence events. Colored branches were tested for positive selection with the branch-site model. Orange bars: p<0.05 for dN/dS>1, red bars: p<0.01 for dN/dS, light blue: p<0.05 for dN/dS<1 and dark blue:p<0.01 for dN/dS<1.

### Cytotoxin-A

*Cytotoxin-A* are cytotoxic proteins that, so far, have only been isolated from Pilidiophora. *Cytotoxin-A* proteins were first isolated from *Cerebratulus lacteus* mucus (Kem and Blumenthal 1978). An homologous toxin was isolated from *Parborlasia corrugatus* mucus and called *Parborlysin* (Berne et al. 2003), both of which were demonstrated to have cytolytic properties. As did Jacobsson et al. (2018), we found a putative transcript for C*ytotoxin-A* in the *Hubrecthella ijimaji* transcriptome, suggesting that such a toxin was present in the ancestor of Pilidiophora. However, these were excluded from further analysis for being too divergent from the remaining species toxins.

### Scoloptoxin SD976-like

We identified scoloptoxins, namely the *Scoloptoxin SSD552* (uncharacterized) and *Scoloptoxin SSD976* (a Voltage-gated calcium channel inhibitor), both of which are cysteine-rich secretory proteins (CRISP) isolated from the chilopoda *Scolopendra subspinipes dehaani* (Liu et al. 2012), in the body transcriptomes and proteomes of *Lineus sanguineus* and *Nemertopsis pamelaroeae* (Tables S3-4). Also, some of the transcripts in the same orthogroup presented similarity to *latisemin*, a CRISP found in the venom of the sea snake *Laticauda semifasciata* (Table S3). However, more experiments are required to determine if the expressed protein is a toxin.

Before filtering, this orthogroup contained many proteins from species of the three classes of Nemertea, and the Annelida *Capitella teleta* (Data not shown), indicating the presence of non-toxin genes in the orthogroup. An hmmscan analysis showed that many of these proteins contain different numbers of ShK-domain like. This domain is similar to the *potassium channel blocker peptide ShK*, described in the sea anemone *Stichodactyla helianthus* (Castañeda et al. 1995). Although this domain has also been described in non-toxins (Möhrlen et al. 2003; Rangaraju et al. 2010), it often plays an important role in controlling potassium channels, as in the *MMP23* (Rangaraju et al. 2010). Moreover, a previous study on Nemertean toxins reported a putative toxin containing the ShK domain (von Reumont et al. 2020), therefore the reported protein likely corresponds to a toxin.

### Alpha-nemertide

The *alpha-nemertides* are peptides that target Na channels with selectivity for arthropods over mammal Na-channels, indicating a biotechnological potential as a pesticide (Jacobsson et al. 2018). The orthology inference showed that these peptides are mostly found in Lineidae, although a transcript was found in *Cerebratulus marginatus*. A thoroughly functional characterization of these toxins’ family has described the effects of different *alpha-nemertides* in Na-channels. However, the role of these peptides in envenomation has yet to be fully elucidated (Jacobsson et al. 2021). Looking closer at the alignment, we can also observe that sequences belonging to different clades differ in the C-Terminal segment, indicating that the product protein may have different properties (Figure S8).

### Beta-Nemertide

The *Beta-nemertide* were described along with the *Alpha-nemertide*; they share some homology with *Neurotoxin-BII*, indicating they may also act as a neurotoxin, but they were not biochemically characterized (Jacobsson et al. 2018). The absence of this toxin sequence from any database complicated its identification, but its low similarity to *Neurotoxins-BII* and its presence in the mucus proteome made us suspect it was a toxin. A pairwise alignment confirmed the identification of the beta-nemertide sequence reported by Jacobsson et al. (2018) (similarity = 94%).

### The Gamma-Nemertides: agelenin-like proteins, a new toxin?

The *Lineus sanguineus* mucus presented peptides of a protein containing a inhibitor cystine knot domain (ICK), a domain present in other known toxins (Gao et al. 2013; Lavergne et al. 2015; Cardoso et al. 2017), including the Alpha-nemertide (Jacobsson et al. 2018). This same protein presents considerable similarity to *Agelenin* (Table S3). *Agelenins* are peptides isolated from the venom of the spider *Agelena opulenta* (Inui et al. 1992), which was demonstrated to produce instantaneous paralysis in cricket (Yamaji et al. 2007). The presence of an ICK domain with similarity to other known neurotoxins, suggests it should be a new neurotoxin, which will be named *Gamma-nemertide*, following the nomenclature of neurotoxins proposed by Jacobsson et al. (2018). Transcripts from *Gamma-nemertide* gene family were found only in the *Lineus* and *Riseriellus* genera.

### alpha-KTx-like/beta-defensin-like/Myticin-like

Lastly, we found an intriguing putative toxin in the mucus proteome, which had the scorpion potassium channel toxin *alpha-KTx 26.2* (e-value=1.7) as the best blast hit and *Beta-defensin* as phammer best hit (e-value=0.0023). Also, hmmscan indicates that it resembles Myticin, an antimicrobial peptide found in mussels (Mitta et al. 1999). Nonetheless, hmmscan also indicates that two transcripts showed low similarity to a domain that confers K+ Channel blocking activity (Zhang et al. 2004), contained in the family of the aforementioned scorpion toxin. This protein shows similarities to both a K+ Channel blocker and a pore-forming protein such as the *Beta-defensin* (van Dijk et al. 2008). In fact, the employment of the defensin domain is recurrent in venoms from other animals, including neurotoxins from scorpions (Fry et al. 2009). Also, we found two different clades of this gene in *Lineus sanguineus*, one of which contains transcripts with longer sequences and motifs somewhat similar to the scorpion toxin domain (Toxin_2; Figure S11). Although we considered this a putative toxin, a non-excluding hypothesis is that it acts as an antimicrobial peptide, forming pores in pathogenic bacteria that may infect the exposed tegument of Nemertea.

Besides being found throughout the Pilidiophora, homologs were also found in transcriptomes from both *Cephalothrix* species examined, but were absent from the remaining Palaeonemertea and Hoplonemertea.

We selected the gene families most likely to be toxins for further analyses. The *cytotoxin-A* and the *alpha-nemertides* had their toxicity well described in previous studies (Kem and Blumenthal 1978; Berne et al. 2003; Jacobsson et al. 2018; Jacobsson et al. 2021). Although no biochemical description of the *beta-nemertides* has been made to our knowledge, its presence in the mucus with similarity to the *Neurotoxin-BII,* which has been biochemically characterized (Kem 1976), made Jacobsson et al. (2018) consider it a toxin. Although toxicity assays are necessary to assert the role of the remaining gene families as toxins, we can still hypothesize their role as toxins based on their similarity to other toxins and their presence in the mucus, a defensive secretion.

### Gene Duplications in Nemertea

The alignment of toxin transcripts to the *L. longissimus* genome allowed to identify the possible genomic location for each toxin gene. For c*ytotoxin-A*, five *loci* were found in tandem (Figure 3), whereas s*coloptoxin SD976-like* transcripts were mapped to six different genomic locations. When mapping only the transcripts retained by MaxAlign, this number decreases to two (Figure S12). For the a*lpha-nemertide*, the number of mapped regions was four, three of which were in the same genome scaffold. The remaining three putative toxin transcripts mapped to two genomic regions each, although some alignments present no intron, indicating potential pseudogenization events (Figure 4; Table S6). These alignments not only revealed the number of copies for these genes in the genome, but also showed that many of these copies are in tandem. Evidence for tandem duplication of toxin genes is also observed in spiders (Bhere et al. 2014) and snakes (Almeida et al. 2021).

**Figure 3:**
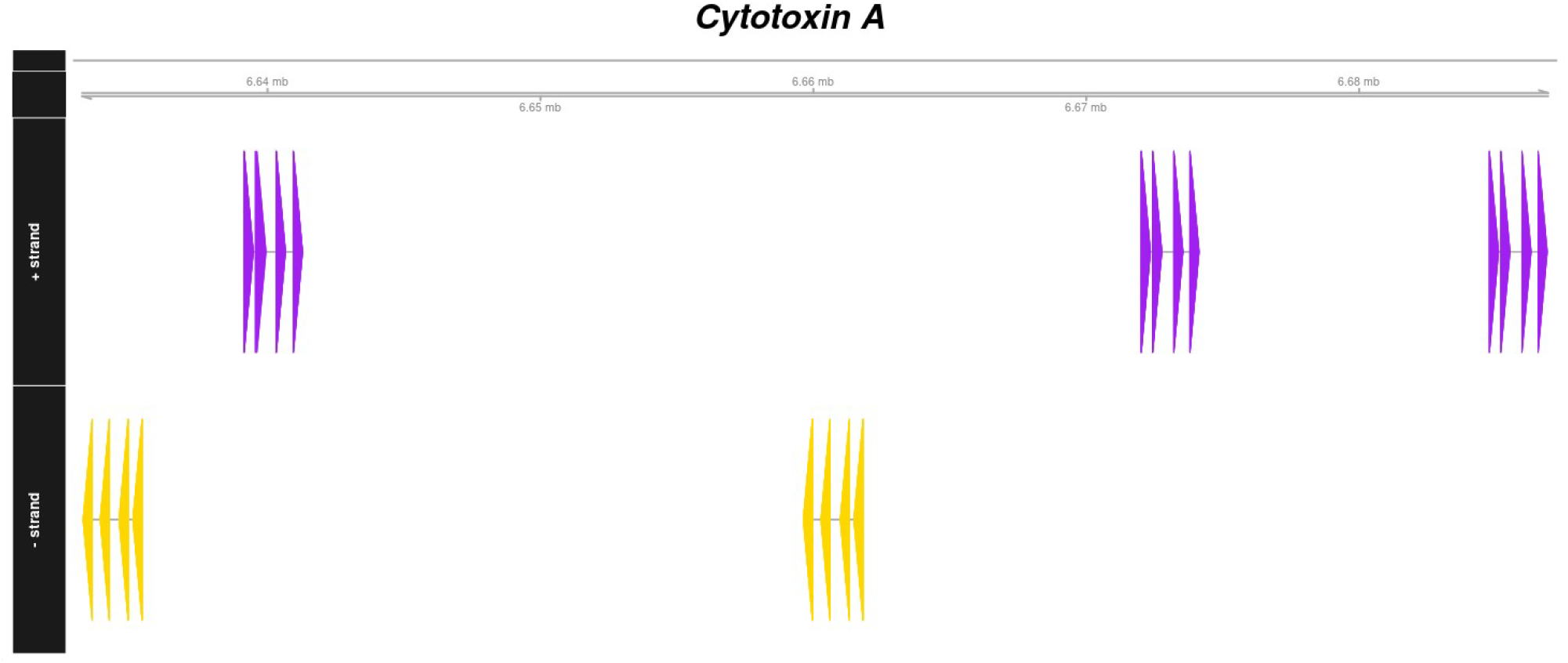
Genomic regions of *Lineus longissimus* aligned to Cytotoxin-A transcripts. Different colors represent the different strands. All the reported alignments covered 100% of the query transcript.

**Figure 4:**
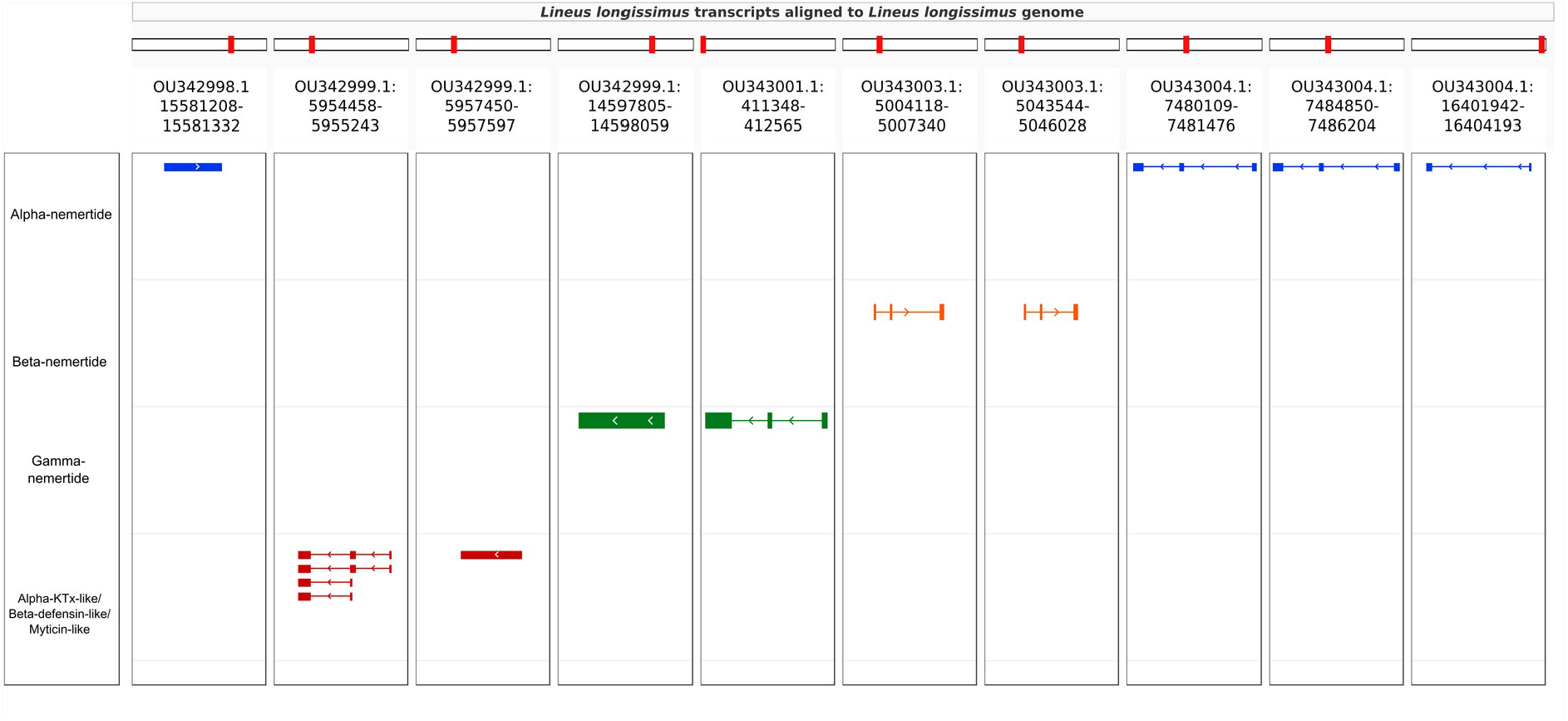
Genomic regions of *Lineus longissimus* containing aligned to the transcripts of the analyzed toxins present in the mucus of *Lineus sanguineus*. The header of each column contains the chromosome name and the range of represented bases. Different colors represent the different toxins prospected.

A tree reconciliation method also provided insights into the number and phylogenomic depth of duplications in toxins in Nemertea (Figures 2, S7-S11; Table S6). By combining this method with the genome alignment, we were able to observe that expansion of the analyzed toxin genes is not a feature restricted to the *Lineus longissimus*, but is spread throughout the Pilidiophora lineage. The presence of transcripts derived from these older duplications in the transcriptome of different species is evidence that these gene copies are present and expressed in many taxa.

The genomic alignments and the reconciled trees results were also shown to be complementary, allowing a deeper understanding of the evolution of toxin gene families. For instance, we assembled four *Cytotoxin-A* transcripts in *Lineus sanguineus*, mapped to five different genomic regions (Figure 3). However, these transcripts belong to only one of the three clades descending from gene duplications in the *Cytotoxin-A* phylogeny (Figure 2), while transcripts from the remaining clades were not found in *L. longissimus*. This could result from a lack of expression from copies belonging to the remaining clades or differences in the number of gene copies between species. For example, in *Notospermus geniculatus,* 11 *Cytotoxin-A loci* were found (Luo et al. 2017), more than twice the number we report in *L. longissimus*.

Duplications have an essential role in the evolution of toxins, either by giving birth to the new genes that can be modified into toxins or by increasing the number of genetic copies of those toxins (Casewell et al. 2013). Multigene families are known for toxin genes of different venomous animals (Duda Jr and Palumbi 1999; Fry et al. 2003; Pi et al. 2006; Duda Jr and Remigio 2008; Juárez et al. 2008; Binford et al. 2009; Chang and Duda Jr 2012; Ma et al. 2012; Zhu et al. 2012; Sunagar et al. 2013; Bayona-Serrano et al. 2020; Giorgianni et al. 2020), and here we show that this seems to be the case for Nemertea as well. New copies may be advantageous at first as they may allow for simultaneous transcription of more than one copy, resulting in augmented gene expression (Casewell et al. 2020). Moreover, it may also facilitate the diversification of the venom by neofunctionalization of one of the copies, which could result in positive selection (Walker 2020). The diversity of targets may determine the fitness change resulting from the diversification of the venom. In snakes, for instance, there seems to be a positive correlation between the complexity of venom and the phylogenetic distance of its preys (Davies and Arbuckle 2019; Lyons et al. 2020; Holding et al. 2021).

Gene duplications leading to venom diversity as an evolutionary response to the number of possible targets could partially explain the increased number of toxin genes of *Lineus sanguineus*, which has a more generalist diet that includes a diversity of polychaetes, oligochaetes and other nemerteans (McDermott and Roe 1985). In contrast, animals from the genus *Nemertopsis* have only been documented to feed on barnacles (Caplins et al. 2012). With such a specialized diet, there would be no selective pressure toward diversifying predatory toxins since the current specialized arsenal efficiently subdues the prey. However, this does not explain the lack of defensive toxins in the mucus of *N. berthalutzae*, since it inhabits the same kind of habitat as *Lineus sanguineus*, and could encounter the same predators. A possible explanation is that, in some groups, non-proteinaceous toxins play a major role in envenomation, whereas peptides are prevalent in others. This could be why the characterized proteinaceous toxins from Nemertea were all isolated from Pilidiophora, while many non-proteinaceous toxins have been isolated and characterized in Hoplonemertea (Göransson et al. 2019).

Similar correlations between the ecology and the venom of *Ototyphlonemertes erneba* can not be clearly established. Although we detected multiple transcripts coding for a single toxin in *O. erneba* (Table S5), discretion is needed before considering these multiple copies of the same toxin, as we sequenced a pool of individuals, meaning the observed diversity may correspond to polymorphisms instead of duplications. Either way, these multiple toxin transcripts indicate diversity either at population or individual level for toxin sequences. Additionally, we identified toxins exclusive to this species, which had not been previously described in Nemertea. We know from rare gut content observations that species of O*totyphlonemertes* may feed on annelids and crustaceans (Corrêa 1948; JLN, pers. obs.), and from Corrêa’s (1949) method for capturing *Ototyphlonemertes* with freshly dead fish that they can be necrofagous. There is, however, no description of the role of venom in prey capture and defense for these meiofaunal animals. Nevertheless, the unusually wide diversity of proboscis structure (Envall and Norenburg 2001) among species of the genus suggests the possibility of strong prey preferences (JLN, pers. obs.). One might expect similar diversity of unique toxins among the species.

Considering the important role of gene duplication in toxin evolution (Kordiš and Gubenšek 2000; Casewell et al. 2013), the finding of more putative toxin genes in both *Lineus sanguineus* transcriptome and proteome, in addition to the fact that proteinaceous toxins have only been isolated from Pilidiophora (Göransson et al. 2019), raises the question if this class has experienced more duplications than the remaining Nemertea. Using CAFE and OrthoFinder, we estimated the number of gene duplication events in each node in the species tree to answer this question. The number of duplications in each node inferred by CAFE ranged from 7 to 3,311 (Figure 5), while in OrthoFinder, this number ranged from 32 to 5,334 (Figure S13).

**Figure 5:**
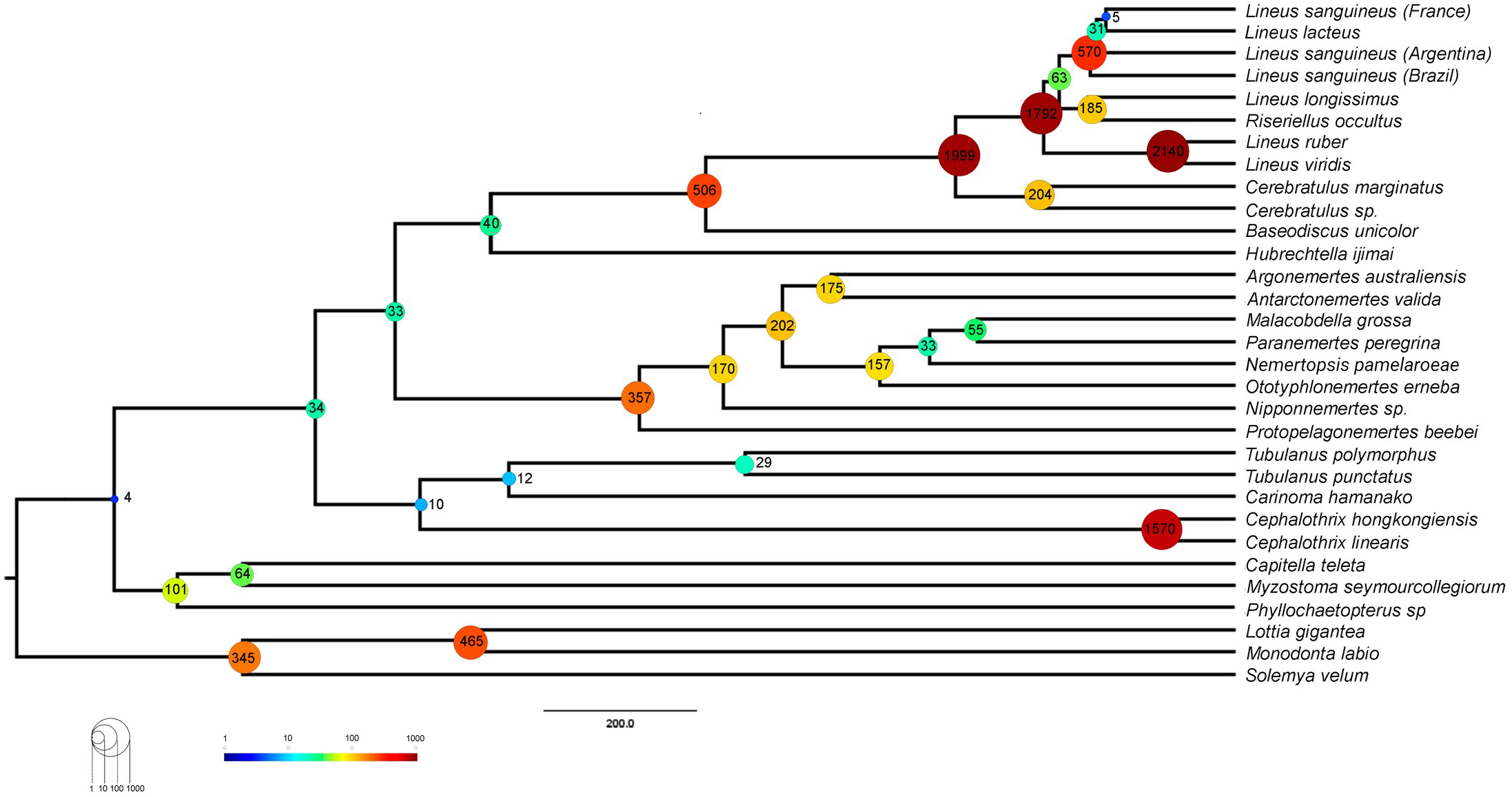
Number of expansions events in each node of the species tree obtained with CAFE analysis allowing for the three different lambdas and including an assembly error model. Numbers are plotted to the ultrametric tree used in CAFE analysis.

CAFE also allowed us to estimate the rate of expansion and contraction events per gene family per Million years (ƛ) in different branches of the tree. Restricting a single value for the whole tree, we found a ƛ of 7,23E-04 (Table 2). In the two ƛ scenario, Pilidiophora had higher ƛ (1,44E-03) than the remaining taxa (7,23E-04), similar to the one in the single ƛ scenario. Adding a third ƛ for the three *Lineus sanguineus* clade did not cause major changes in the other two ƛ. Adding an error model to deal with assembly errors did not change the ƛ drastically for Pilidiophora or the remaining taxa, but slightly changed the ƛ for *Lineus sanguineus*. All of the scenarios allowing more than one lambda were significantly more likely than the global lambda scenario (P<0.01) (Table 2).

Our results (Figure 5, Figure S13) show that nodes within Pilidiophora have more duplications than the remaining Nemertea. Consistent with these results, estimates of ƛ were significantly higher in Pilidiophora when compared to the remaining groups, even corrected for the multiple sampling of *Lineus sanguineus* and accounted for assembly errors using CAFE errors model (Table 2). Gene duplications result in redundant proteins, which may experience neofunctionalization and lead to evolutionary novelties (Lewis 1951; Qian and Zhang 2014; Birchler and Yang 2022). Alternatively, as discussed earlier, new copies allow for simultaneous transcription, potentially increasing the expression of the duplicated gene (Casewell et al. 2020). Either way, such an increased rate of gene duplications in Pilidiophora could have resulted in genetic novelties, such as the origin of new toxins or the expansion of toxin gene families.

Besides Pilidiophora, the palaeonemertean *Cephalothrix* also displays an increased number of gene duplications, when compared to the remaining Nemertea. This, coupled with the findings of Whelan et al. (2014), which described a high diversity of toxins in the genus, may indicate that this group is a good candidate for future proteinaceous toxin prospection. Interestingly, species from the *Cephalothrix* were also found to have tetrodotoxin in their body, a strong toxin produced by symbiotic bacteria (Magarlamov et al. 2014).

### Selection test in toxins

To better understand how the sequence of these toxins evolved, we applied selection tests using models accounting for codons with heterogeneous evolution rates. In the alignments of all six analyzed genes, the presence of sites under positive selection was statistically significant under the site-model (p<0.05), and at least a single codon under positive selection was found in each gene family (Table 3). Moreover, the branch-site model, which additionally allows heterogeneity between branches of the gene tree was applied for the *Cytotoxin-A*, s*coloptoxin SD976-like* and a*lpha-nemertide* gene families. C*ytotoxin-A* and *alpha-nemertide* had one branch presenting sites under positive selection (Table 3). Toxin sequence alignment and the sites under positive selection can be found on Figures 2, and S6-S10.

As the main driver of selection in toxin genes is the toxin target (the prey or predator that will be affected by such a toxin), the substitution in these positively selected sites could lead to changes in the toxin activity and specificity as an evolutionary response to the different selective pressures imposed by the ecological challenges faced by each species. In other words, these amino acid substitutions could be an evolutionary response to the changes in the composition of its targets. Many of these sites under selection fell within toxin domains, supporting this hypothesis (Figures S6, S9 and S10).

For all tested toxins, except for *Scoloptoxin SD976-like*, we found evidence of positive selection within clades delimited by duplications (branch-site model) or without duplications (site model). This indicates that natural selection led to toxin divergence between different species. Put in another way, changes in these sites in toxin genes may be being driven by a selective pressure for the toxin to work better against the targets of each species. Positive selection between species has been previously described in toxins (Rokyta et al. 2011; Zhu et al. 2016).

At the same time, we observe both gene duplications and sites under positive selection for the *Cytotoxin-A*, *Scoloptoxin SD976-like* proteins, *alpha-nemertides* and b*eta-nemertides*. For the C*ytotoxin-A* gene family, the site detected under positive selection by the site-model was not the same detected by the branch-site model. The diversity in sites found to be under positive selection only by the site model could be explained by natural selection favoring the divergence of paralogs toxins. Gene duplications with positive selection between the copies lead to an increase in venom complexity, which may allow for an increase in the diversity of potential molecular targets of these toxins by neofunctionalization. Diversifying potential molecular targets may help overcome eventual resistances evolved in the target. Alternatively, it may allow for the inclusion of new species in the diet, or act as a defense mechanism against new potential predators. Such positive selection after duplications has been previously described in toxins (Chang and Duda Jr 2012; Zhang et al. 2015; Modahl et al. 2018) and physiological mammal genes (Han et al. 2009).

To test how toxin evolution is related to the natural history of Nemertea is a challenging task. As our data show, different species might employ the same toxin in different contexts (*i.e.,* defense and predation). Also, the natural history of ribbon worms needs to be better elucidated due to their secretive way of life, thus the knowledge of actual prey and predators for different species of Nemertea is scarce.

Either way, these hypotheses leading to target diversification and specialization are not only testable, but also interesting to guide future searches for toxins that may have pharmacological applications. The expected small changes in the toxin caused by these sites under positive selection may be of interest to the development of pharmaceuticals and pesticides, as they are likely a result of selective pressure for changes in the toxin activity and selectivity.

## Final considerations

In this work, we prospected for putative toxins in three Nemertea species using a proteo-transcriptomic approach. We report a high diversity of putative toxin genes in *Lineus sanguineus*, with orthologs in other species, but a not-so-diverse arsenal in the analyzed hoplonemerteans genera *Nemertopsis* and *Ototyphlonemertes*. To the best of our knowledge, this is the first study to systematically assess the molecular evolution of toxin genes in Nemertea, accounting for gene duplications and selection tests. We show that the toxin genes observed for these animals evolve similarly to those described in snakes, spiders, scorpions and cone-snails. As in these groups, Nemertea toxin genes may experience positive selection and gene duplications. Nevertheless, there is still much to be elucidated on the biochemical characteristics of these toxins and their role of these toxins in the natural history of ribbon worms. Summing up, our study and other recent publications on the topic reveal an extensive diversity of toxins produced by these animals, which up until now have been relatively neglected. Beyond any doubt, there is a lot to learn from these curious animals, and what we know so far is but a small piece of this intriguing puzzle: the toxins produced by Nemertea.

## Acknowledgments

We would like to thank Instituto de Biociências – USP, the Faculdade de Farmácia – USP and the Centro de Biologia Marinha da Universidade de São Paulo (CEBIMar) staff; all the colleagues from Laboratório de Diversidade Genomica, especially Cecili Mendes, Tammy Arai, Dione Jordan and everyone else who assisted in fieldwork. Additionally, we are grateful to Darwin server administrators for their assistance, to the Food Research center (FAPESP, Grant 2013/07914-8) staff for their help with the proteomic analyses. We also would like to thank Håkan Andersson for discussing the mucus extraction protocol with us and Rafael Iwama for helping with the analyses of anticoagulant motifs in antistasin. This work was funded by CNPQ and the Fundação de Amparo à Pesquisa do Estado de São Paulo (FAPESP, grants 2015/20139-9, 2018/12502-4, and 2020/06467-1). The authors declare that they have no known competing financial interests or personal relationships that could have appeared to influence the work reported in this paper.

## Author contributions

GGS and SCSA conceived the project. First version of the manuscript was written by GGS and SCSA. GGS, JLN and SCSA collected the specimens. Material preparation, data collection and analysis were performed by GGS, JPF and ECT. All authors contributed to the manuscript writing and approved the submitted version.

## Supplementary Figures

Figure S1: Analyses pipeline. Dataset 1 was used to infer a well supported species tree and estimate overall gene duplication rates. Dataset 2 was used to infer toxin orthogroups, from which gene duplications analyses and selection tests were performed. Dataset 3 was used to search for proteins in the mucus and body proteomes from the analyzed species, in order to identify and validate putative toxins. aa= amino acid.

Figure S2: BUSCO assessment results. The used database was the odb9 for metazoans conserved genes (n=978 genes), hence each hit corresponds to a conserved metazoan gene.

Figure S3: Occupancy matrix for the 5,208 genes used in the phylogenomic step. Dark tiles represent orthogroups present. Species are ordered from highest to the lowest occupancy.

Figure S4: Venn diagram of putative toxin best hits found in the transcriptome of the three species sequenced in this study.

Figure S5: Abundance of toxins in the proteomic sample, indirectly measured by the summed area of peptide peaks in the HPLC chromatogram, calculated by PEAKS Studio Xpro, for mucus and body fractions replicates from a) *Lineus sanguineus* and b) *Nemertopsis berthalutzae*. Ar: Body sample of *L. sanguineus* collected in Araçá; P: Body sample of *L. sanguineus* collected in Peró; Nem: Body sample of *N. berthalutzae* collected in Praia Grande; 1: Fraction of proteins with 50 KDa or more; 2: Fraction of proteins between 20KDa and 50KDa; 3: Fraction of proteins of 20KDa or lower; IDs between parentheses correspond to IDs from tables S3 and S4.

Figure S6: Protein sequence coverage by HPLC-MS/MS of *Antistasin-like protein*. Peptides found are represented by light blue lines. Detected amino-acid modifications are indicated by boxes.

Figure S7: Gene tree reconciliation, alignment and omega value for sites of the *Scoloptoxin SDD976-like* curated gene family. Legend goes as in Figure 2. ShK: ShK domain-like.

Figure S8: Gene tree reconciliation, alignment and omega value for sites of the *alpha-nemertide* gene family. Legend goes as in Figure 2.

Figure S9: Gene tree reconciliation, alignment and omega value for sites of the B*eta-nemertide* gene family. Legend goes as in Figure 2.

Figure S10: Gene tree reconciliation, alignment and omega value for sites of the *Gamma-nemertide* gene family. Legend goes as in Figure 2. Toxin_35: Toxin with inhibitor cystine knot ICK or Knottin scaffold.

Figure S11: Gene tree reconciliation, alignment and omega value for sites of the *alpha-KTx/Beta-defensin/myticin* gene family. Legend goes as in Figure 2. YiaAB: yiaA/B two helix domain; Toxin_2: Scorpion short toxin, BmKK2; Myticin-prepro:Myticin pre-proprotein from the mussel.

Figure S12: Genomic region containing regions identified as homologous to the *Scoloptoxin-SD976-like* putative toxins. The only *Lineus longissimus* transcript included in the final *Scoloptoxin-SD976-like* alignment was the LlongTRINITY_DN3733. OU342995.1: Scaffold name from *Lineus longissimus* genome assembly.

Figure S13: Gene duplications per node identified with OrthoFinder and duplication support value higher than 50%. A support of at least 50% means that at least half of the species present in given node is also present at the two direct children nodes.

## Supplementary Tables

Table S1: OTUs used for phylogeny construction and selection test and their respective accession code. Outgroup samples are marked with asterisks.

Table S2: Statistics of the assembled transcriptomes. RIN= RNA integrity number.

Table S3: Annotation of the predicted putative toxins of Lineus sanguineus. Y: yes; S: Successfully predicted.

Table S4: Annotation of the predicted putative toxins of *Nemertopsis pamelaroeae.* Y: yes; S: Successfully predicted.

Table S5: Annotation of the predicted putative toxins of Ototyphlonemertes erneba.

Table S6: Number of gene copies resulting from duplication events inferred from tree reconciliation method. Genome alignment are the loci number of loci inferred from alignments to *Lineus longissimus* genome for each orthogroup.

