## Supplementar Figures and Tables for "Venomous Noodles: evolution of toxins in Nemertea through positive selection and gene duplication"

### Supplementar file

#### Supplementar Figures:

- Figure S1
- Figure S2
- Figure S3
- Figure S4
- Figure S5
- Figure S6
- Figure S7
- Figure S8
- Figure S9
- Figure S10
- Figure S11
- Figure S12
- Figure S13

#### Supplementar tables

- Table S1
- Table S2
- Table S3
- Table S4
- Table S5
- Table S6

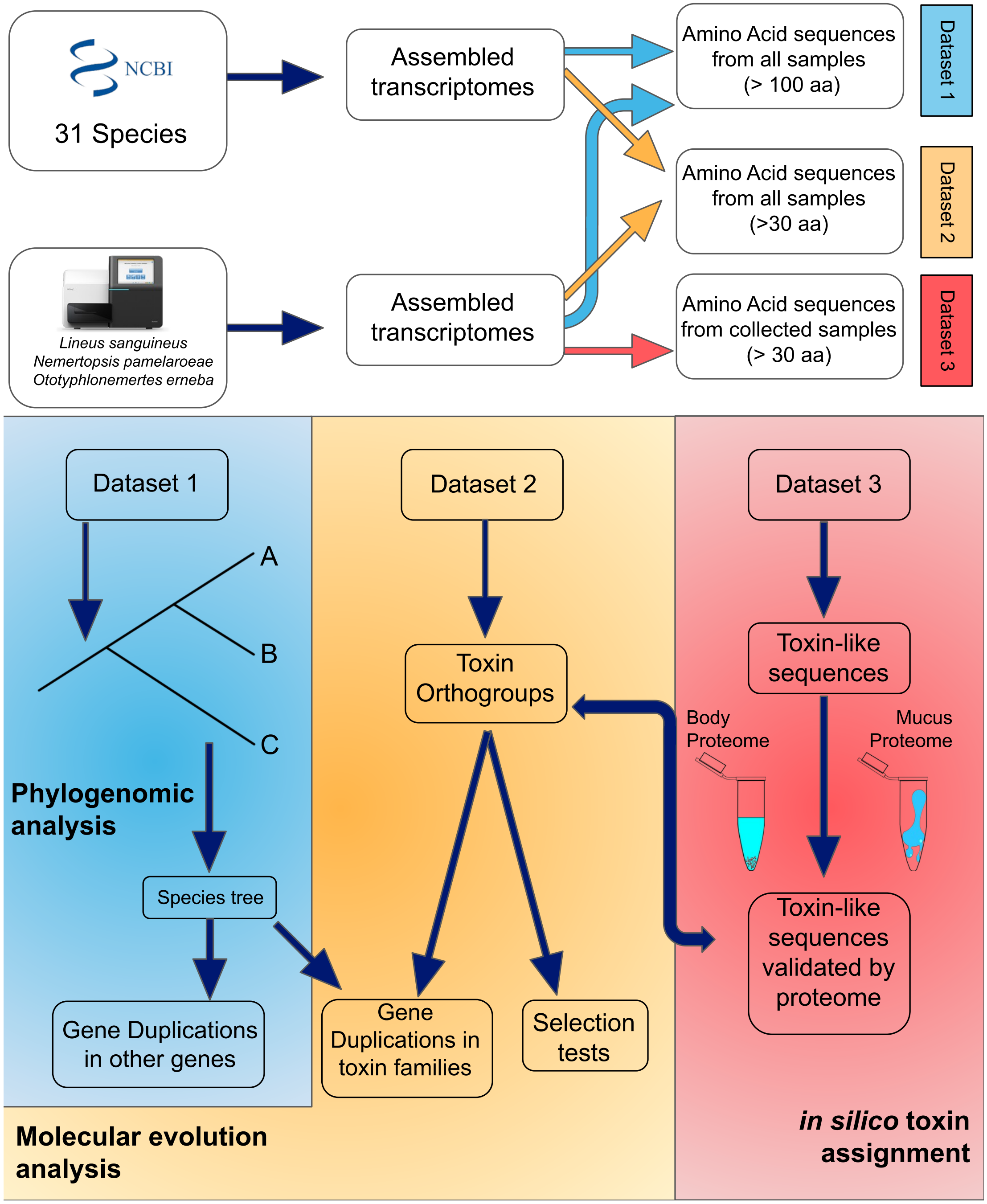


**Figure S1:** Analyses pipeline. Dataset 1 was used to infer a well supported species tree and estimate overall gene duplication rates. Dataset 2 was used to infer toxin orthogroups, from which gene duplications analyses and selection tests were performed. Dataset 3 was used to search for proteins in the mucus and body proteomes from the analyzed species, in order to identify and validate putative toxins. aa= amino acid.

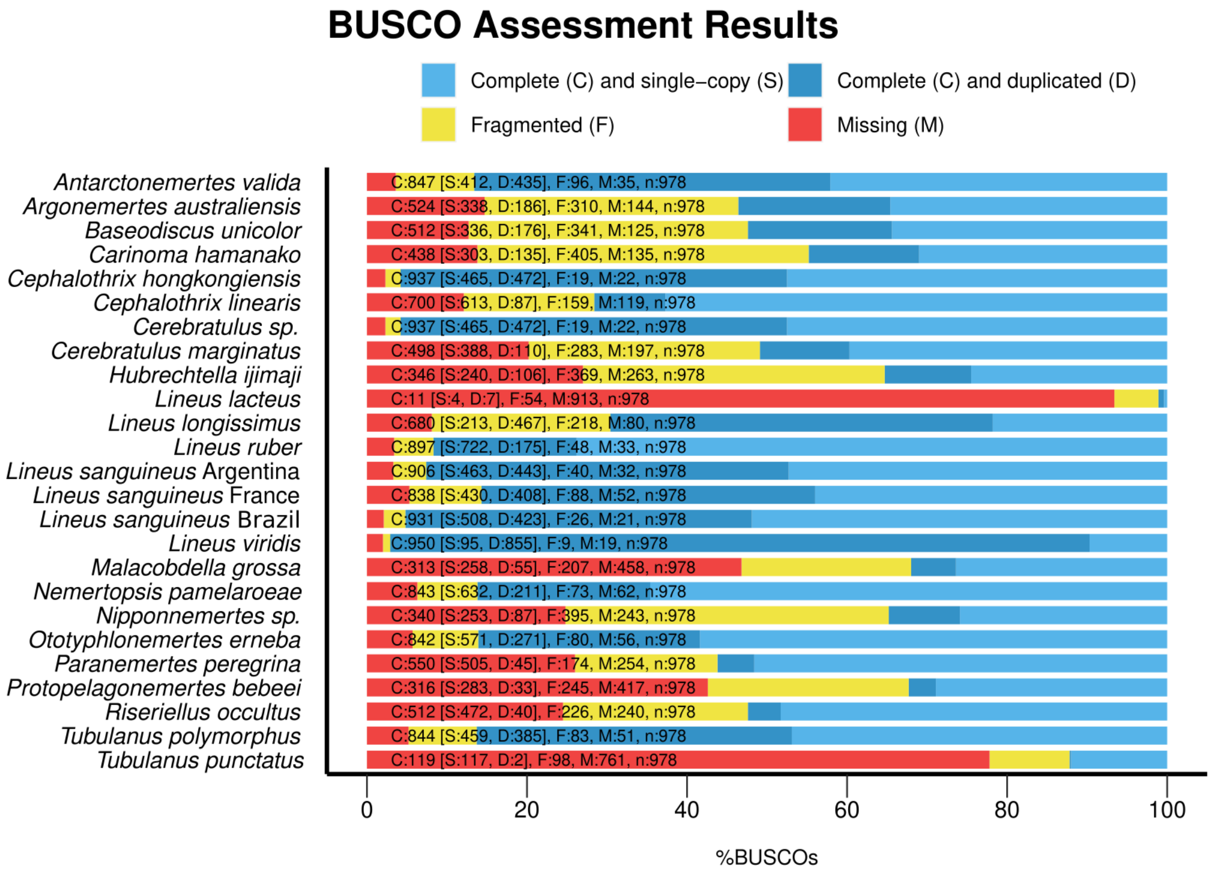


**Figure S2:** BUSCO assessment results. The used database was the odb9 for metazoans conserved genes (n=978 genes), hence each hit corresponds to a conserved metazoan gene.

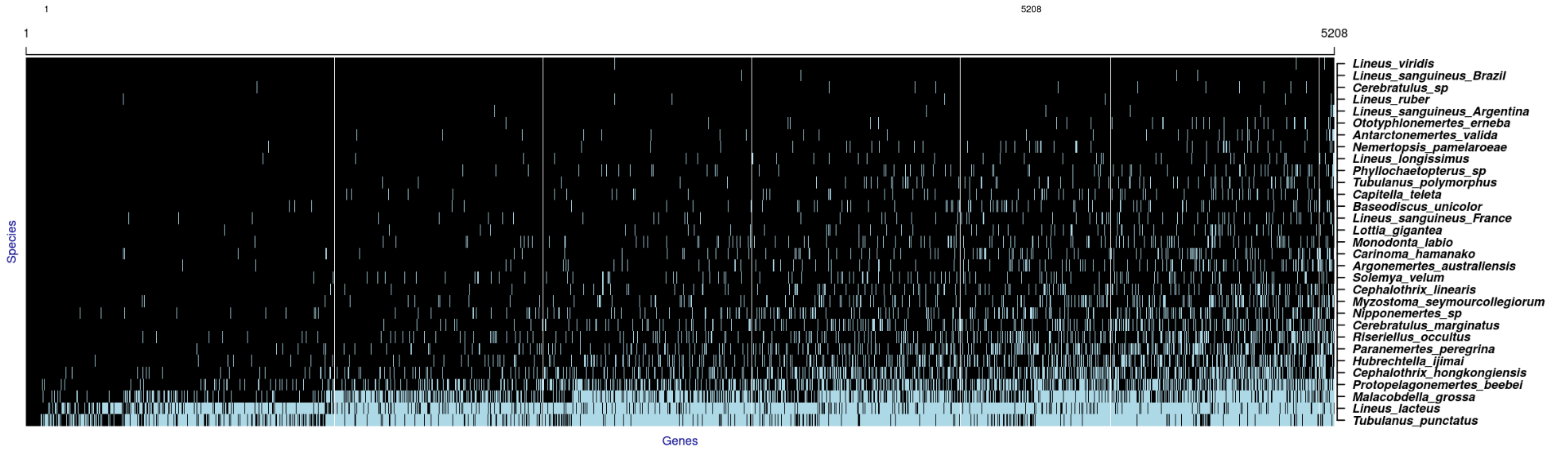


**Figure S3:** Occupancy matrix for the 5,208 genes used in the phylogenomic step. Dark tiles represent orthogroups present. Species are ordered from highest to the lowest occupancy.

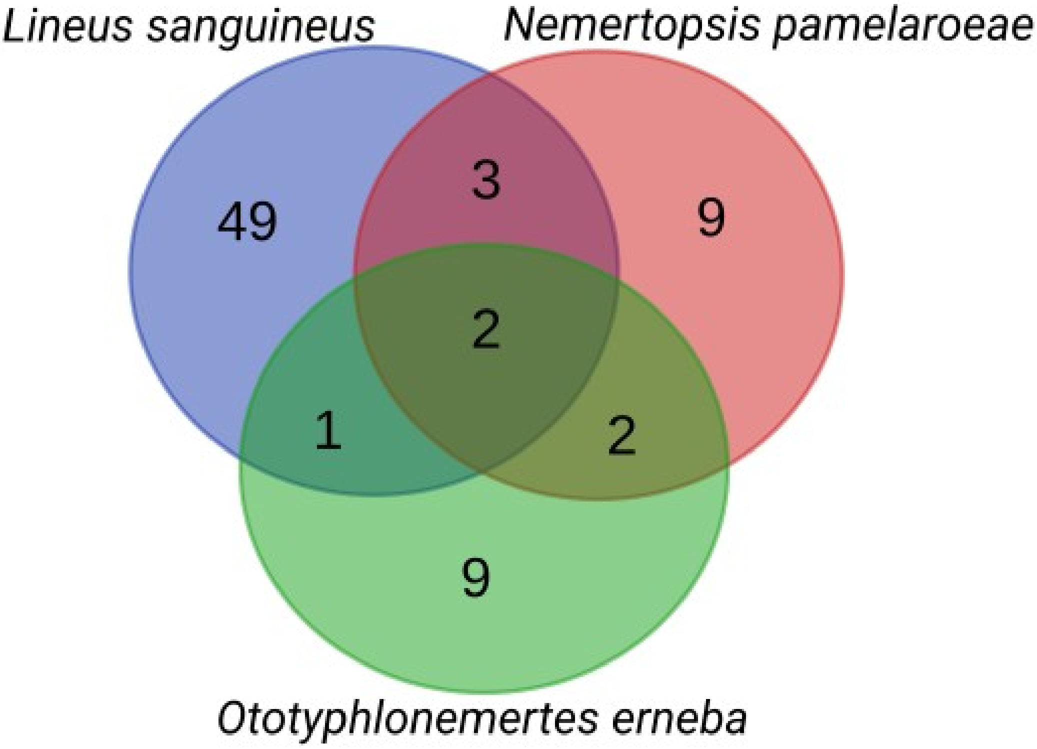


**Figure S4:** Venn diagram of putative toxin best hits found in the transcriptome of the three species sequenced in this study.

a)

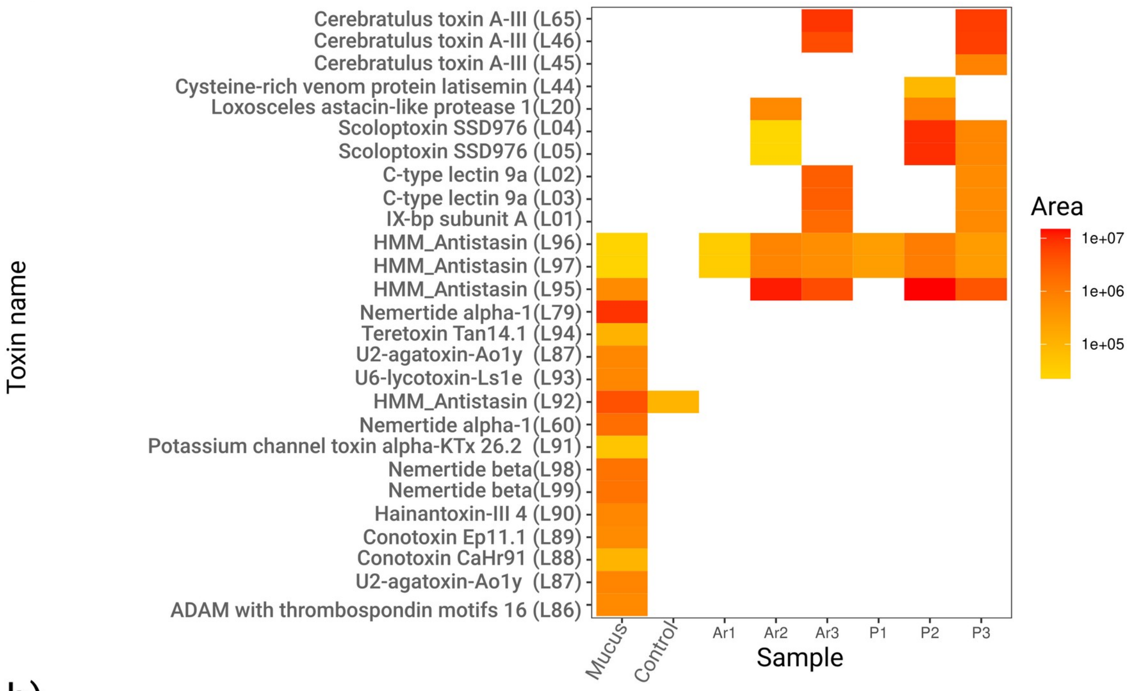


b)

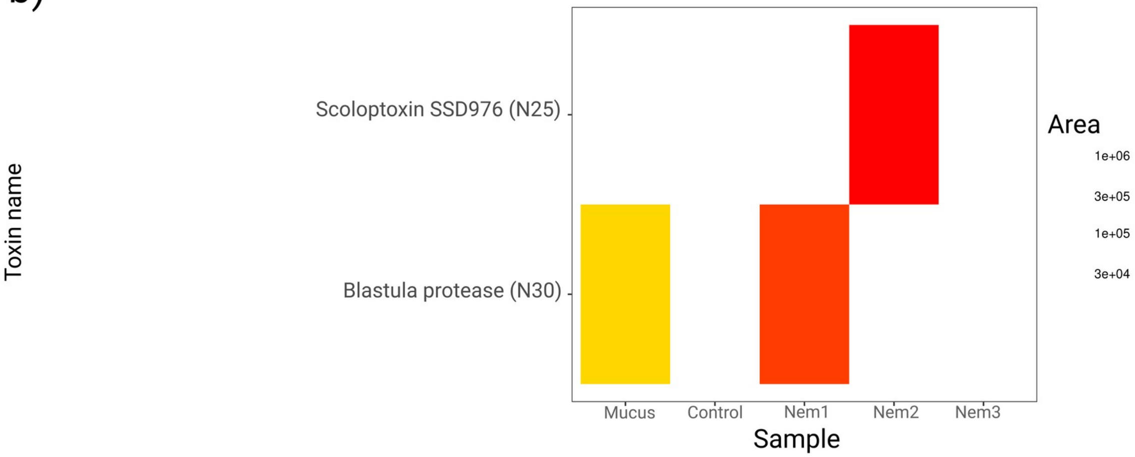


**Figure S5:** Abundance of toxins in the proteomic sample, indirectly measured by the summed area of peptide peaks in the HPLC chromatogram, calculated by PEAKS Studio Xpro, for mucus and body fractions replicates from a) *Lineus sanguineus* and b) *Nemertopsis berthalutzae*. Ar: Body sample of *L. sanguineus* collected in Araçá; P: Body sample of *L. sanguineus* collected in Peró; Nem: Body sample of *N. berthalutzae* collected in Praia Grande; 1: Fraction of proteins with 50 KDa or more; 2: Fraction of proteins between 20KDa and 50KDa; 3: Fraction of proteins of 20KDa or lower; IDs between parentheses correspond to IDs from tables S3 and S4.

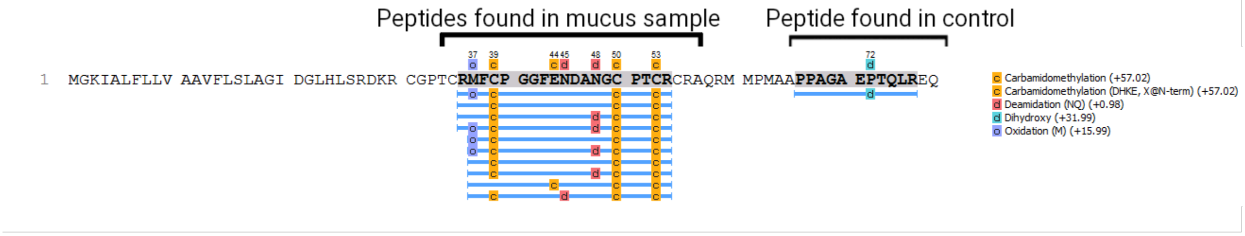


**Figure S6:** Antistasin (L92) protein sequence coverage by HPLC-MS/MS of *Antistasin-like protein*. Peptides found are represented by light blue lines. Detected amino-acid modifications are indicated by boxes.

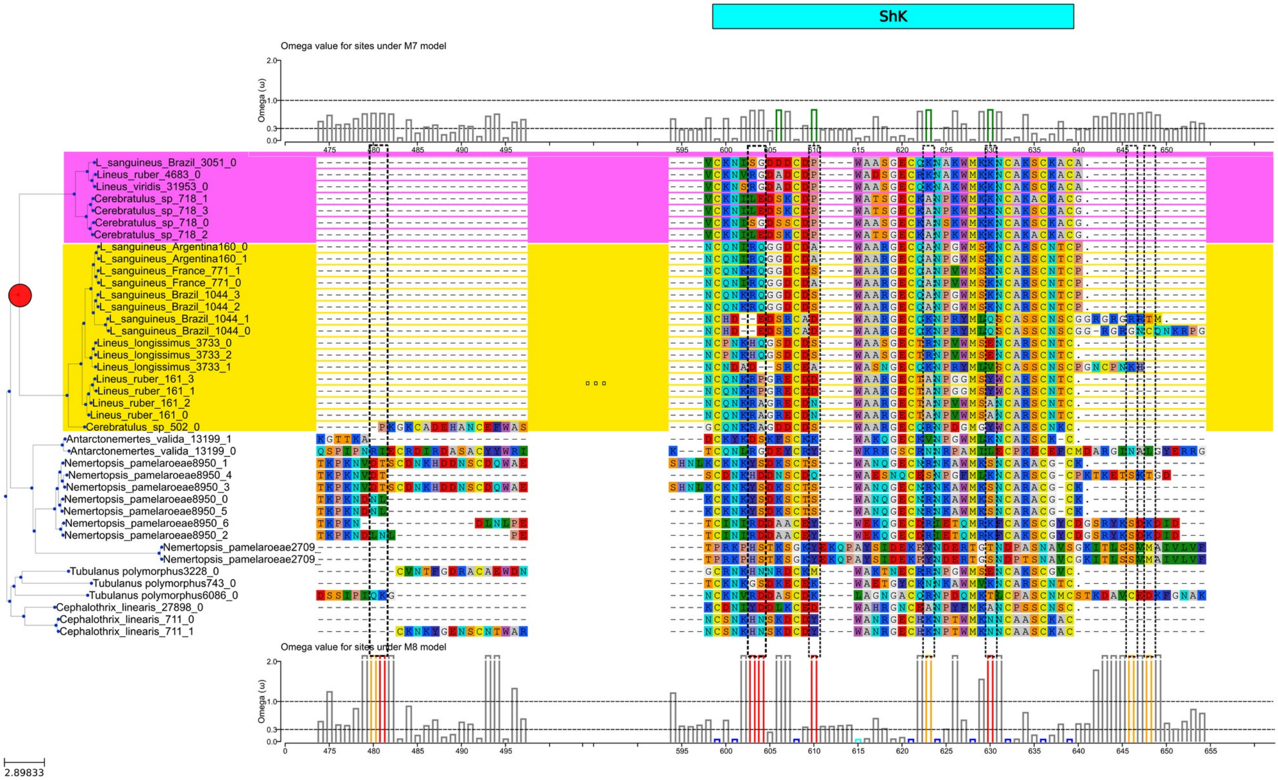


**Figure S7:** Gene tree reconciliation, alignment and omega value for sites of the *Scoloptoxin SDD976-like* curated gene family. Legend goes as in Figure 2. ShK: ShK domain-like.

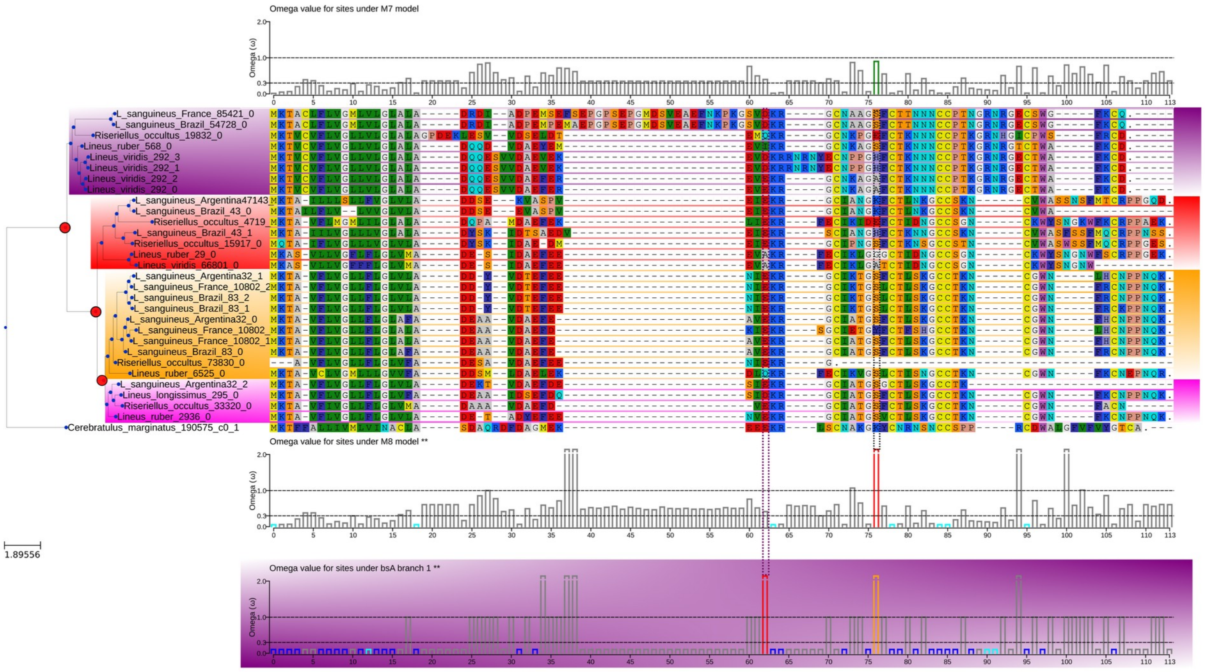


**Figure S8:** Gene tree reconciliation, alignment and omega value for sites of the *Alpha-Nemertide* gene family. Legend goes as in Figure 2.

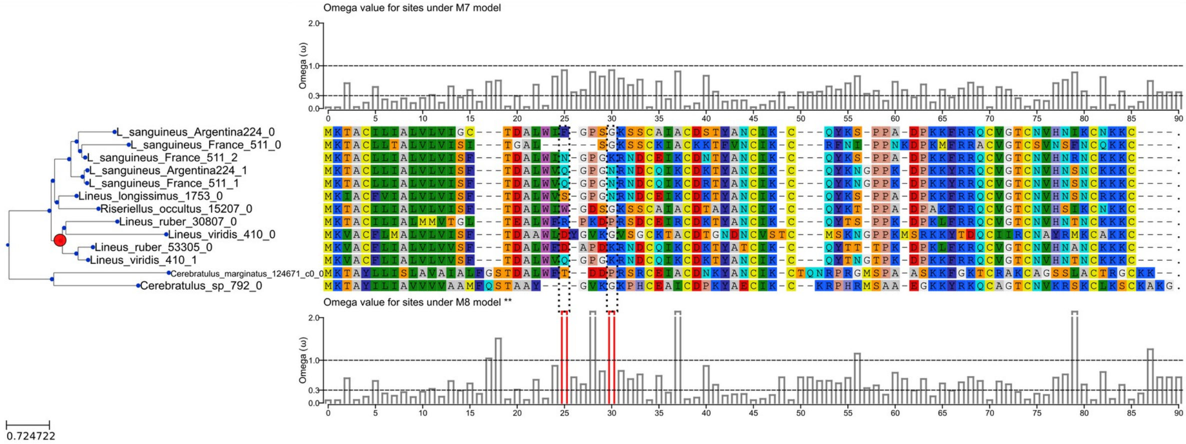


**Figure S9:** Gene tree reconciliation, alignment and omega value for sites of the *Beta-Nemertide* gene family. Legend goes as in Figure 2.

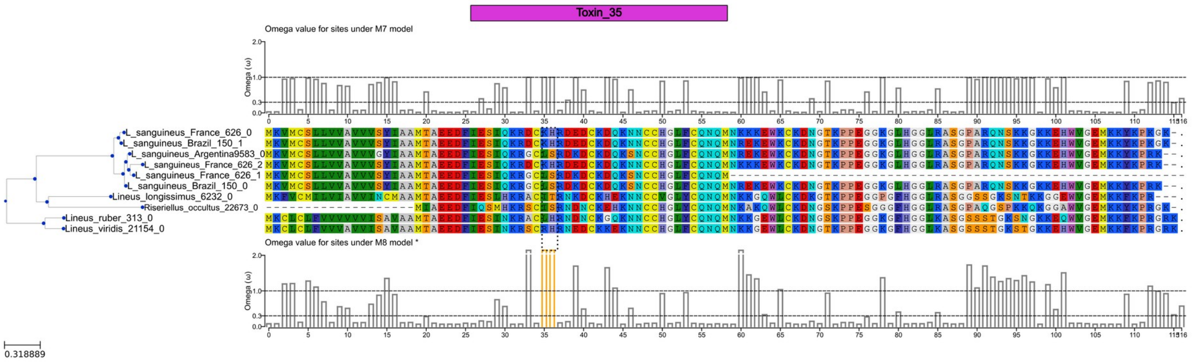


**Figure S10:** Gene tree reconciliation, alignment and omega value for sites of the *Gamma-Nemertide* gene family. Legend goes as in Figure 2. Toxin_35: Toxin with inhibitor cystine knot ICK or Knottin scaffold.

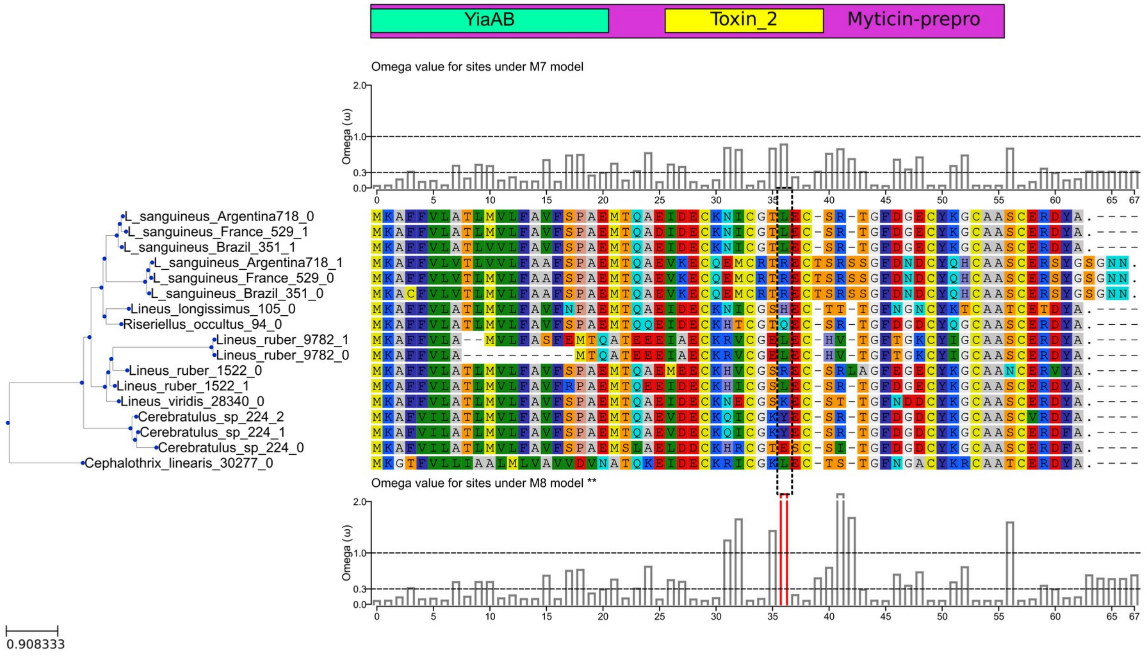


**Figure S11:** Gene tree reconciliation, alignment and omega value for sites of the *alpha-KTx/Beta-defensin/myticin* gene family. Legend goes as in Figure 2. YiaAB: yiaA/B two helix domain; Toxin_2: Scorpion short toxin, BmKK2; Myticin-prepro:Myticin pre-proprotein from the mussel.

# OU342995.1

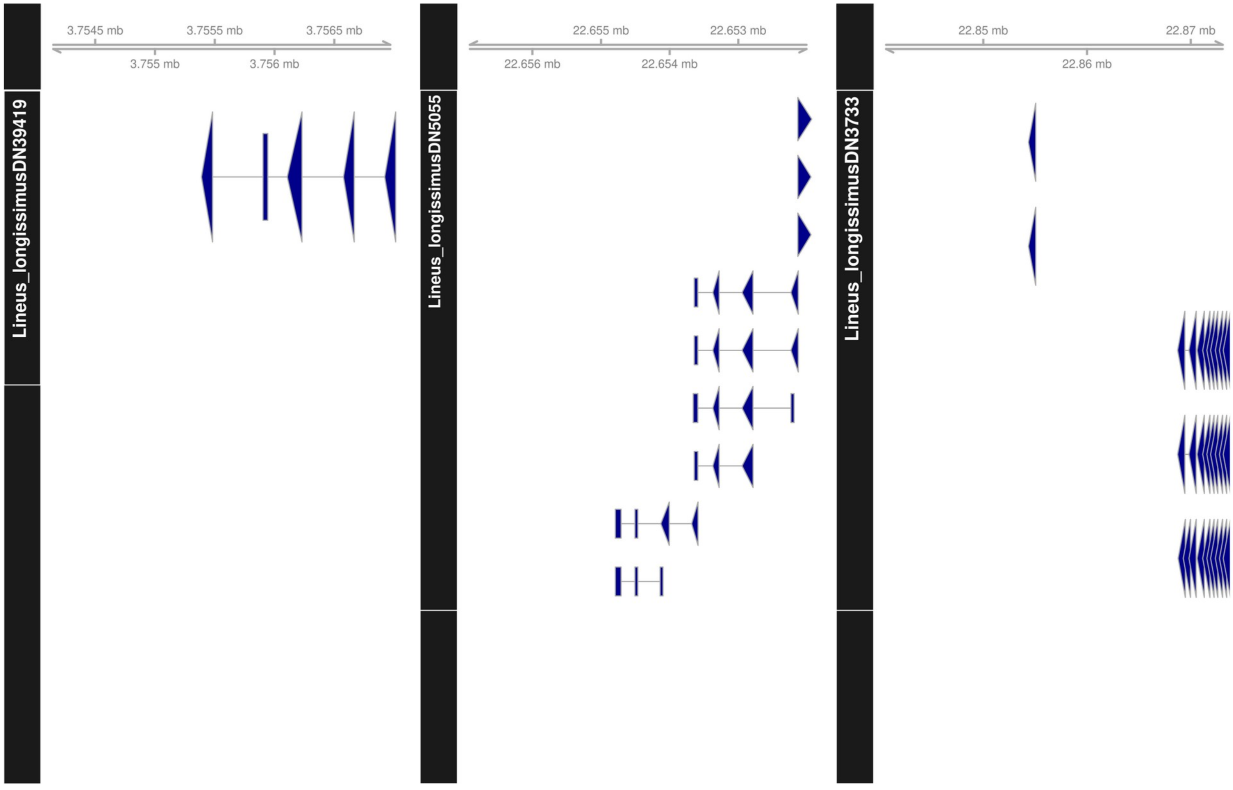


**Figure S12:** Genomic region containing regions identified as homologous to the Scoloptoxin-SD976-like putative toxins. The only Lineus longissimus transcript included in the final Scoloptoxin-SD976-like alignment was the LlongTRINITY_DN3733. OU342995.1: Scaffold name from Lineus longissimus genome assembly.

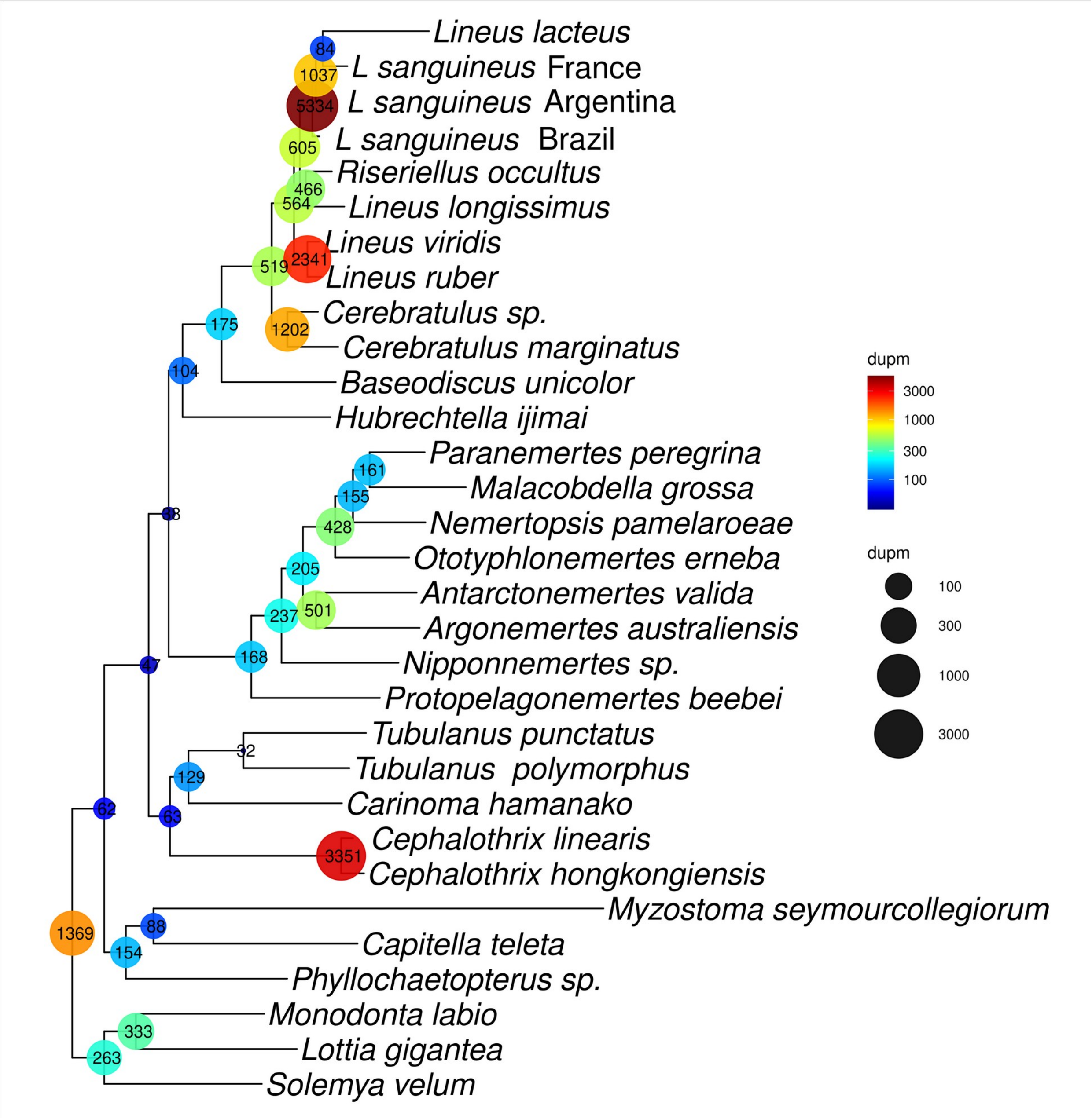


**Figure S13:** Gene duplications per node identified with OrthoFinder and duplication support value higher than 50%. A support of at least 50% means that at least half of the species present in given node is also present at the two direct children nodes.

Table S1: OTUs used for phylogeny construction and selection test and their respective accession code. Ougroup samples are marked with asterisks. Samples from the present work are in bold.

| OTU | SRR, SRX or GCA (When specified) |
|---|---|
| *Antarctonemertes valida* | PRJNA485632 |
| *Argonemertes australiensis* | PRJNA254358 |
| *Baseodiscus unicolor* | PRJNA322119 |
| *Carinoma hamanako* | PRJNA254071 |
| *Cephalothrix hongkongiensis* | PRJNA181263 |
| *Cephalothrix linearis* | PRJNA245790 |
| *Cerebratulus marginatus* | PRJNA181261 |
| *Cerebratulus sp.* | PRJNA275078 |
| *Hubrechtella ijimai* | PRJNA254167 |
| *Lineus lacteus* | PRJNA530965 |
| *Lineus longissimus* | PRJNA330781 |
| *Lineus ruber* | PRJNA249058 |
| *Lineus sanguineus* (Argentina*)* | PRJNA322119 |
| ***Lineus sanguineus (Brazil, BR)*** | **PRJNA952238** |
| *Lineus sanguineus* (France) | PRJNA322119 |
| *Lineus viridis* | PRJNA322119 |
| *Monodonta labio* | PRJNA253054 |
| ***Nemertopsis pamelaroeae*** | **PRJNA952238** |
| *Nipponnemertes sp.* | PRJNA254364 |
| ***Ototyphlonemertes erneba*** | **PRJNA952238** |
| *Paranemertes peregrina* | PRJNA254365 |
| *Protopelagonemertes beebei* | PRJNA254366 |
| *Riseriellus occultus* | PRJNA254175 |
| *Tubulanus polymorphus* | PRJNA263418 |
| *Tubulanus punctatus* | PRJNA254065 |
| *Capitella teleta** | Genome https://metazoa.ensembl.org/Capitella_teleta/Info/Index |
| *Lottia gigantea** | Genome http://metazoa.ensembl.org/Lottia_gigantea/Info/Index |
| *Monodonta labio** | PRJNA253054 |
| *Myzostoma seymourcollegiorum** | PRJNA282832 |
| *Phyllochaetopterus sp** | PRJNA243848 |
| *Solemya velum** | PRJNA72139 |

Table S2: Statistics of the assembled transcriptomes. RIN: RNA integrity number.

| Species | RIN | Number of raw reads | Number of paired filtered reads | #contigs | #contigs >500pb | #contigs >1000pb | longest contig | N50 |
|---|---|---|---|---|---|---|---|---|
| *Lineus sanguineus* | 7.70 | 13.441.397 | 12.229.480 | 113.957 | 65.600 | 44.455 | 30.915 | 3.167 |
| *Ototyphlonemertes erneba* | 7.30 | 12.859.411 | 11.688.148 | 99.617 | 46.875 | 28.177 | 34.319 | 2.417 |
| *Nemertopsis pamelaroeae* | 8.60 | 12.106.267 | 11.008.779 | 54.385 | 26.684 | 15.019 | 19.971 | 2.079 |

Tables S3-S5 are available in TableS3_S5.xls

Table S6:Number of gene copies resulting from duplication events inferred from Tree reconciliation method and number of Loci inferred from alignments to *Lineus longissimus* genome for each orthogroup.

| Orthogroup | Tree reconciliation Method | Genome Alignment |
|---|---|---|
| Cytotoxin A | 3 | 5 |
| Scoloptoxin SD976-like | 2 | 2 * |
| Alpha Nemertide | 4 | 4 |
| Beta Nemertide | 2 | 2 |
| Gamma Nemertide | 1 | 2 |
| KTX-like | 1 | 2 |

*Only the transcripts kept by MaxAlign were aligned to the genome
